# A Benchmark of Semi-Supervised scRNA-seq Integration Methods in Real-World Scenarios

**DOI:** 10.1101/2025.08.23.671952

**Authors:** Xiaoyu Shen, Chuan He, Leying Guan

## Abstract

Semi-supervised methods for single-cell RNA-seq integration promise to improve batch correction and biological signal preservation by leveraging cell-type labels. However, their reported benefits often rely on overly idealized settings. Here, we present the first systematic benchmark of five leading semi-supervised methods (scANVI, scGEN, ssSTACAS, scDREAMER, ItClust) against five widely used unsupervised baselines across six diverse datasets. We evaluate performance under five realistic annotation scenarios, including missing, erroneous, boundary-missing and mixed, batch-specific, and auto-generated labels, using nine established integration metrics. While semi-supervised methods show gains with perfect annotations, their robustness declines sharply under practical imperfections. Only scANVI and ssSTACAS maintain stable but modest improvements relative to their unsupervised counterparts, while none consistently outperform the strongest unsupervised method, scCRAFT. Our results highlight that current semi-supervised strategies offer limited practical advantage and that careful choice of integration method remains critical when label quality is uncertain.

## 1 Introduction

Single-cell RNA sequencing (scRNA-seq) has revolutionized our understanding of cellular diversity and resulted in large-scale resources that capture cellular states under various biological conditions and individuals [1–3]. Initiatives such as the Human Cell Atlas [4] and the Human BioMolecular Atlas Program [5], which aim to map human tissues at unprecedented resolution, have provided valuable insights into cellular heterogeneity in health and disease. However, these large-scale scRNA-seq resources are often affected by batch effects–unwanted technical and biological variation introduced by differences in sample handling, sequencing platforms, protocols, or inter-individual factors such as microenvironmental context [6–9]. To address this challenge, a wide range of computational integration methods have been developed to mitigate batch-specific transcriptomic shifts. Most integration methods are unsupervised and rely solely on the count matrices of the scRNA-seq data and their performance has been extensively benchmarked [10].

Semi-supervised integration methods leverage available cell-type labels to improve embedding quality and batch correction by guiding the alignment process with partial supervision. For example, scANVI [11] and scDREAMER [12] encourage accurate classification of annotated cells to their labels during integration. scGEN [13] models gene expression changes associated with discrete covariates such as batch by learning latent shifts across conditions, which are estimated from matched or re-sampled cell-type distributions to isolate the effect of interest. ItClust [14] applies transfer learning to propagate information from labeled reference cells to unlabeled target cells. More recently, ssSTACAS [15] refines the mutual nearest neighbor anchoring process by pruning cell pairs whose labels are incompatible, thereby improving alignment fidelity. Although improvement of semi-supervised approaches have been observed in their respective studies, their evaluations often rely on idealized scenarios with complete or randomly perturbed labels, failing to reflect real-world challenges where annotations are incomplete, inconsistent, erroneous, or automated with variable accuracy [16–18]. Consequently, the true utility and robustness of these semi-supervised methods in practical applications remain poorly understood.

To bridge this gap, we present a comprehensive benchmark of semi-supervised single-cell RNA-sequencing integration methods under realistic conditions. We systematically evaluate five leading semi-supervised methods (scANVI, scGEN, ssSTACAS, scDREAMER, and ItClust) against five unsupervised methods (Seurat RPCA [19], scVI [20], Harmony [21], Scanorama [22], and scCRAFT[23]) that are selected based on popularity and reported performance (see Supplementary Note 1 for detailed description of each method). Our benchmark design incorporates two commonly examined conditions: (1) randomly missing labels and (2) randomly wrong labels, as well as three realistic scenarios that mimic practical challenges: (3) ambiguous labels at cluster boundaries, (4) batch-specific partial annotations, and (5) labels generated by automated annotation tools.

By assessing semi-supervised and unsupervised methods in different annotation settings using six diverse datasets spanning multiple tissues, organs, and species, 2 our study provides the first systematic and practically grounded evaluation of semi-supervised integration strategies, offering critical guidance for their effective use in real-world single-cell analyses.

## 2 Results

### 2.1 Benchmark Design Overview

This study considers in total six annotation scenarios. The first is a baseline setting where unsupervised methods perform integration, and supervised methods are provided with correct labels for all cells. The other five are partial label scenarios, two of which have been widely used in previous work to test the robustness of semi-supervised methods: one with randomly missing labels and another with randomly incorrect labels, each at varying degrees. (Fig. 1).

**Fig. 1.**
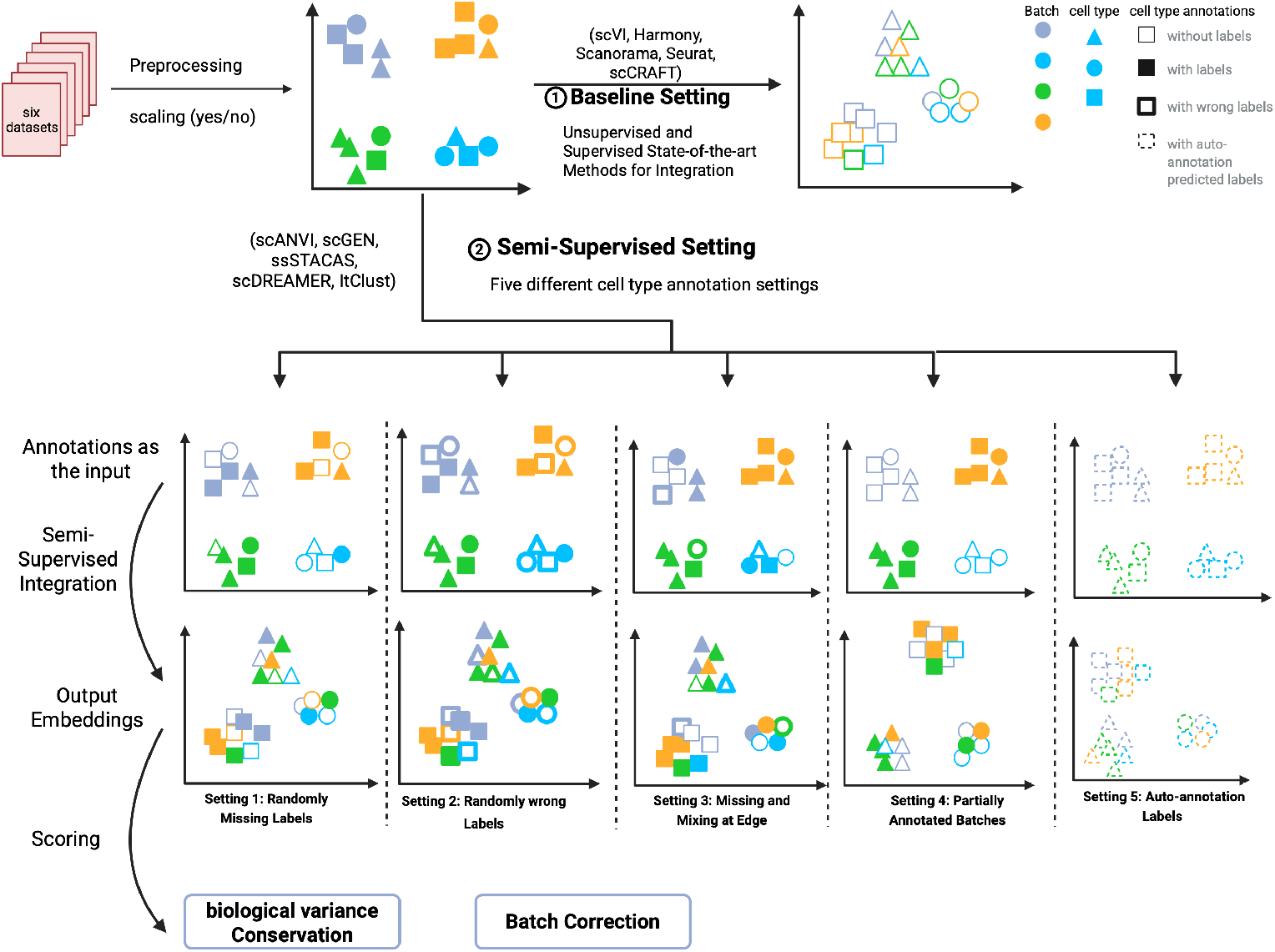
Design of semi-supervised integration benchmarking. Schematic diagram of the benchmarking workflow. In this study, ten data integration methods, including five semi-supervised algorithms and five unsupervised baselines, are evaluated across six integration datasets. Five label scenarios are designed to reflect diverse real-world conditions, including randomly missing, randomly wrong, missing and mixing at edge, partially annotated batches, and automatically generated labels, together with the baseline setting where five unsupervised methods are used and five semi-supervised method are used with full labels presented. Integration results are assessed using 9 metrics that evaluate batch effect removal and conservation of biological variance from true cell type labels (label conservation).

The third partial label scenario is *missing and mixing at edge*. A key factor in achieving strong biological conservation is the separation of such closely related cell types where even a small number of correctly annotated cells can help reinforce this distinction and improve integration performance [10]. However, misannotations in practice often arise specifically at these boundaries [23]. To account for this, we introduces the scenario where labels are mixed between transcriptionally similar cell types, rather than randomizing labels arbitrarily. Concretely, in the *missing and mixing at edge* scenario, we first conduct unsupervised integration with Harmony, then, we examine the 30 nearest neighbors for each cell and compute the proportion of neighbors that share its true cell type. If this proportion falls below a threshold *γ ∈* [0, 1], then the cell’s label is reassigned to a different type sampled from its neighbors’ label distribution. If the resampled label still matches the cell’s true type, we instead mark the label as “unknown”. Thus, a higher *γ* value selects more cells for label alteration. Each selected cell’s label is then changed to a neighboring cell type or “unknown.” Consequently, increasing *γ* results in more significant label corruption and a more difficult annotation task.

The fourth partial label scenario, *partially annotated batches*, simulates situations where only a randomly selected subset of batches is labeled. This reflects a common challenge in practice. For example, when integrating datasets from multiple studies, researchers can rely on author-provided manual annotations. However, such annotations are not always available for all studies, and differences in annotation granularity across atlases can introduce inconsistencies [24, 25]. In some cases, it is common to manually annotate or select a representative subset of datasets or batches to create partial high-quality labels for the full atlas [26, 27]. The *partially annotated batches* scenario is thus designed to evaluate how well integration methods perform under incomplete batch-wise supervision.

Finally, the fifth partial label scenario, *auto-annotation labels*, evaluates the use of labels generated by three popular automated annotation tools, which are often only partially correct. The increasing availability of single cell atlases and collective efforts to define cell types based on prior knowledge are valuable information for cell-type identifications [14]. Auto-annotation tools can generate annotations based on this knowledge [16–18, 28–30]. Although inherently imperfect, auto-annotations offer a practical and scalable alternative to labor-intensive manual curation. This scenario allows us to assess how well semi-supervised integration methods perform when provided with machine-generated labels, an increasingly relevant setting in large-scale single-cell analysis.

We evaluate different methods across the above scenarios using six datasets, namely, the human pancreas, mammary epithelial cell (bct), macaque, human immune, lung atlas, and lung two species datasets, which vary in the number of cells, batches, and cell populations (cell types), in order to represent different levels of challenges in the scRNA integration task. We use the average overall performance of unsupervised methods to determine the relative difficulty of each datasets. Among the six datasets, 4 the human pancreas dataset [31] is the easiest for unsupervised integration, containing 16,382 cells across 9 batches. This is followed by the mammary epithelial cell (bct) dataset [32–34], with 9,288 cells and 3 batches, and the macaque dataset [35], which includes 30,302 cells from 30 batches. The most challenging data set is the lung two species dataset [36], which comprises of 2 batches from human and mouse lung cells and introduces a significant cross-species heterogeneity. The second most difficult is the lung atlas dataset [37], with 32,472 cells across 16 batches, followed by the human immune dataset [38], which contains 33,506 cells from 10 batches. Detailed preprocessing procedures and datasets descriptions are provided in Methods and Supplementary Note 2, respectively.

Each integration method was evaluated with nine metrics grouped into two categories: (i) conservation of biological variation and (ii) removal of batch effects (Fig. 1). Biological conservation was quantified with global cluster-matching metrics—Adjusted Rand Index (ARI) [39] and normalized mutual information (NMI) [40]—together with silhouette width by cell-type label (ASW label) and cell-type LISI (cLISI) [21]. Batch-effect removal was assessed using silhouette width by batch (ASW batch) [41], the k-nearest-neighbor batch-effect test acceptance rate (kBET) [41], batch LISI (bLISI) [42], k-nearest-neighbor graph connectivity, and the true positive rate (TPR) [43]. Although TPR reflects both biological conservation and batch mixing, it is empirically more sensitive to batch mixing; following recent practice [23], we therefore treat TPR as a batch-removal metric in our summaries. Overall accuracy scores were computed as a weighted mean of the two category averages, with 60% weight on biological conservation and 40% on batch-effect removal, following [10] (see Methods).

### 2.2 Baseline Setting with Oracle Labels

We compare five semi-supervised integration methods—scANVI, scGEN, ssSTACAS, scDREAMER, and ItClust using oracle labels, defined as the original cell-type annotations provided with each dataset. These methods are evaluated both against each other and against five unsupervised approaches—Seurat RPCA, scVI, Harmony, Scanorama, and scCRAFT to assess their label-utilization efficiency under ideal labeling conditions. In addition, the performance of the unsupervised methods serves as a baseline to reflect the intrinsic integration difficulty of the six datasets. Based on the average overall performance of these unsupervised methods, the datasets are ranked from easiest to most difficult as follows (with average unsupervised methods’ overall score listed in parenthesis): human pancreas (0.7519), bct (mammary epithelial cells) (0.7496), macaque (0.7249), human immune (0.6825), lung atlas (0.6773), and lung two-species (0.6329) (Fig. 2). scDREAMER generally achieved the highest overall weighted score among all semi-supervised methods, outperforming the strongest unsupervised competitor, scCRAFT, by an average of 7.05% across the six datasets (Fig. 2a-b). Notably, scDREAMER surpassed all other semi-supervised and unsupervised integration methods across all six datasets in terms of batch removal, outperforming the second-best method, scCRAFT, by 12.18% after averaging across all datasets (Fig. 2a-c). In contrast, ItClust excelled in biological conservation performance, outperforming the second-best method, scGEN, by 7.28% after averaging across all datasets.

**Fig. 2.**
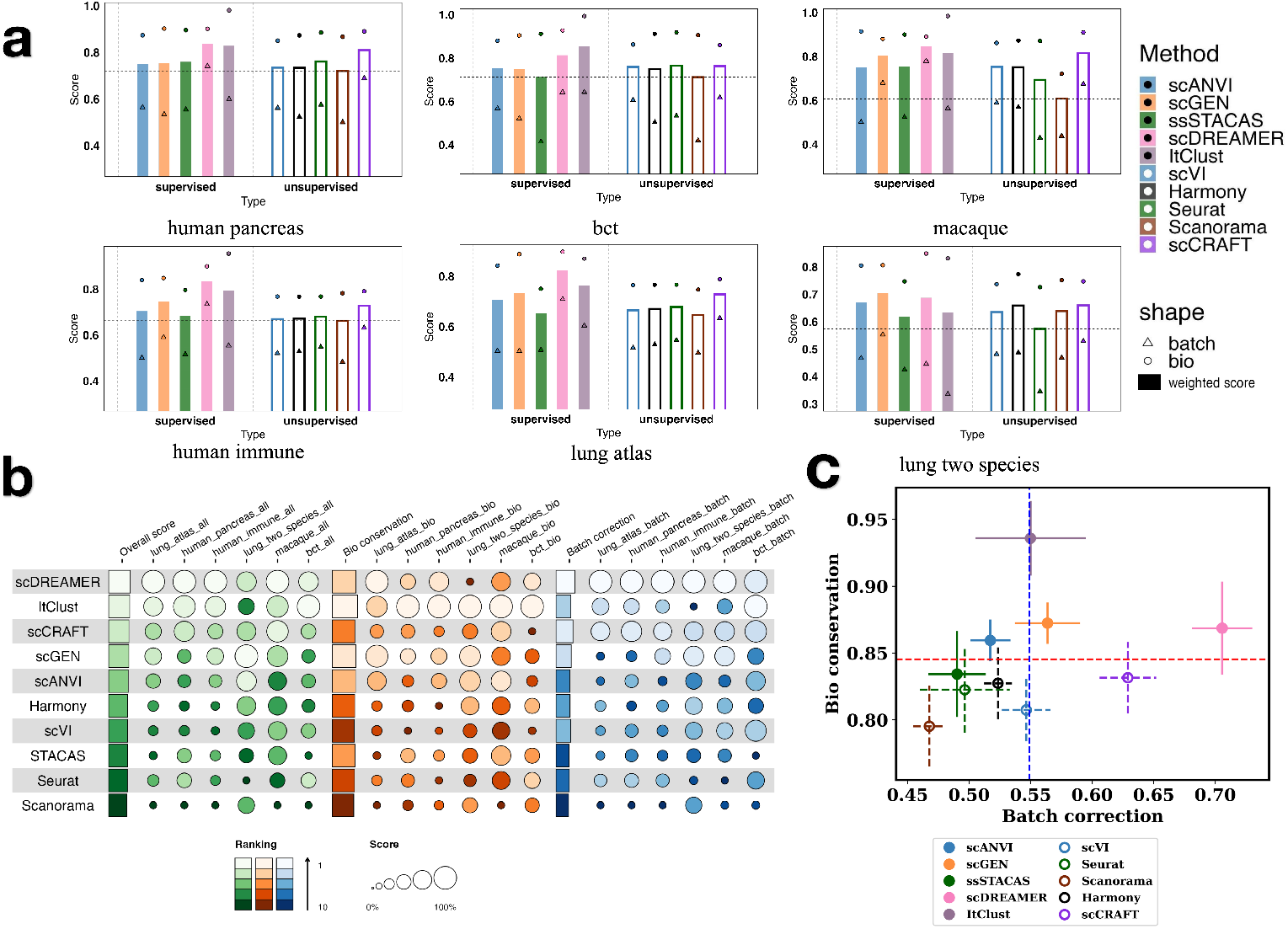
Baseline settings: using unsupervised methods results. (a) Bar plots showing the performance of all methods across six datasets under this setting. Each bar represents the overall weighted score of a method; triangles and circles indicate the batch correction and biological conservation scores, respectively. In each dataset sub-figure, the left panel displays the performance of five semi-supervised methods provided with full labels, represented by solid-colored bars. The right panel shows the performance of five unsupervised methods, depicted with empty (unfilled) bars. (**b**) Bubble plot of all the methods relative ranking considering the overall weighted score, bio conservation score and batch correction score. The first column of each color group represents the average performance of each method across all six datasets. (**c**) Scatter plot of the average overall batch correction score against the average overall bio conservation score for each methods across all datasets. The error bars represent the standard error of the mean for each method and score type across all datasets. The horizontal red dashed line represents the average bio conservation score across all methods, while the vertical blue dotted line represented the average batch correction score.

In this ideal baseline for semi-supervised integration, the other three semi-supervised approaches also generally attain higher overall weighted scores than their unsupervised counterparts when averaged across all datasets, with robust improvement in terms of biological conservation but varied performance in batch removal (Fig. 2a-c). Specifically, scANVI and scGEN both perform better than scVI in the biological conservation task, with average improvements of 6.45% and 8.04% respectively. However, considering the batch removal, while scGEN is 3.17% higher than scVI, scANVI performs worse than scVI by 5.63% on average. ssSTACAS, whose structure is similar to Seurat RPCA, shows overall similar performance to it, performing better at the macaque and lung two-species datasets but slightly worse in other four datasets. It showed a slight improvement in overall and biological conservation performance, but slightly lower batch correction performance.

In summary, although semi-supervised approaches tend to improve biological conservation relative to their unsupervised integration backbones, their overall performance and advantage vary substantially when compared to other state-of-the unsupervised approaches. Only scDREAMER, ItClust, and scGEN performed at least as well as unsupervised methods in both biological conservation and batch removal, while showing clear improvements in at least one of these aspects (Fig. 2c). When compared to scCRAFT, the strongest unsupervised method, ssSTACAS performed worse in both biological conservation and batch removal. Among the other semi-supervised approaches, although varying degrees of improvement in biological conservation were observed, only scDREAMER emerged as a clear winner. This is primarily due to substantially lower batch removal scores observed for ItClust, scGEN, and scANVI. These observations highlight the importance of employing suitable and efficient modeling strategies for both transcriptomic variation and label information in achieving successful single-cell RNA-seq integration.

### 2.3 Scenarios I and II: Randomly Missing or Wrong Labels

In Scenario I, we evaluated the robustness of five semi-supervised integration methods to missing annotations by randomly removing 30%, 50%, and 70% of cell-type labels in each dataset. This scenario has been widely examined previously, where semi-supervised approaches often outperform their unsupervised counterparts.

ssSTACAS and scANVI, while less effective at leveraging correct labels compared to scDREAMER, ItClust, and scGEN in the oracle setting, were robust to randomly missing labels, showing minimal performance deterioration even at 70% missingness. Both methods performed better than their respective unsupervised backbones (Seurat RPCA and scVI, respectively). In contrast, ItClust exhibited the highest sensitivity to missing labels, showing the largest drop in bio-conservation scores with merely 30% randomly missing labels and performing worse than all other unsupervised or semi-supervised approaches on every dataset except for *bct*. At 70% missingness, it consistently underperformed across all datasets (Fig. 3a). scGEN and scDREAMER also showed substantial declines in performance compared to the baseline oracle setting, with deterioration increasing alongside missingness. scANVI remained the top-performing semi-supervised integration method at 70% missingness, outperforming the second-best, ssSTACAS, by an average of 3.78% in overall score across all datasets primarily due to higher bio-conservation scores, but still 3.85% lower compared the best unsupervised method, scCRAFT. However, a closer look reveals that scDREAMER achieved the best or near-best performance across most datasets, except for macaque (which has 30 batches) and lung two species (which has two batches). These exceptions contributed to scDREAMER’s 5.82% lower overall score compared to scANVI. (Fig. 3a).

**Fig. 3.**
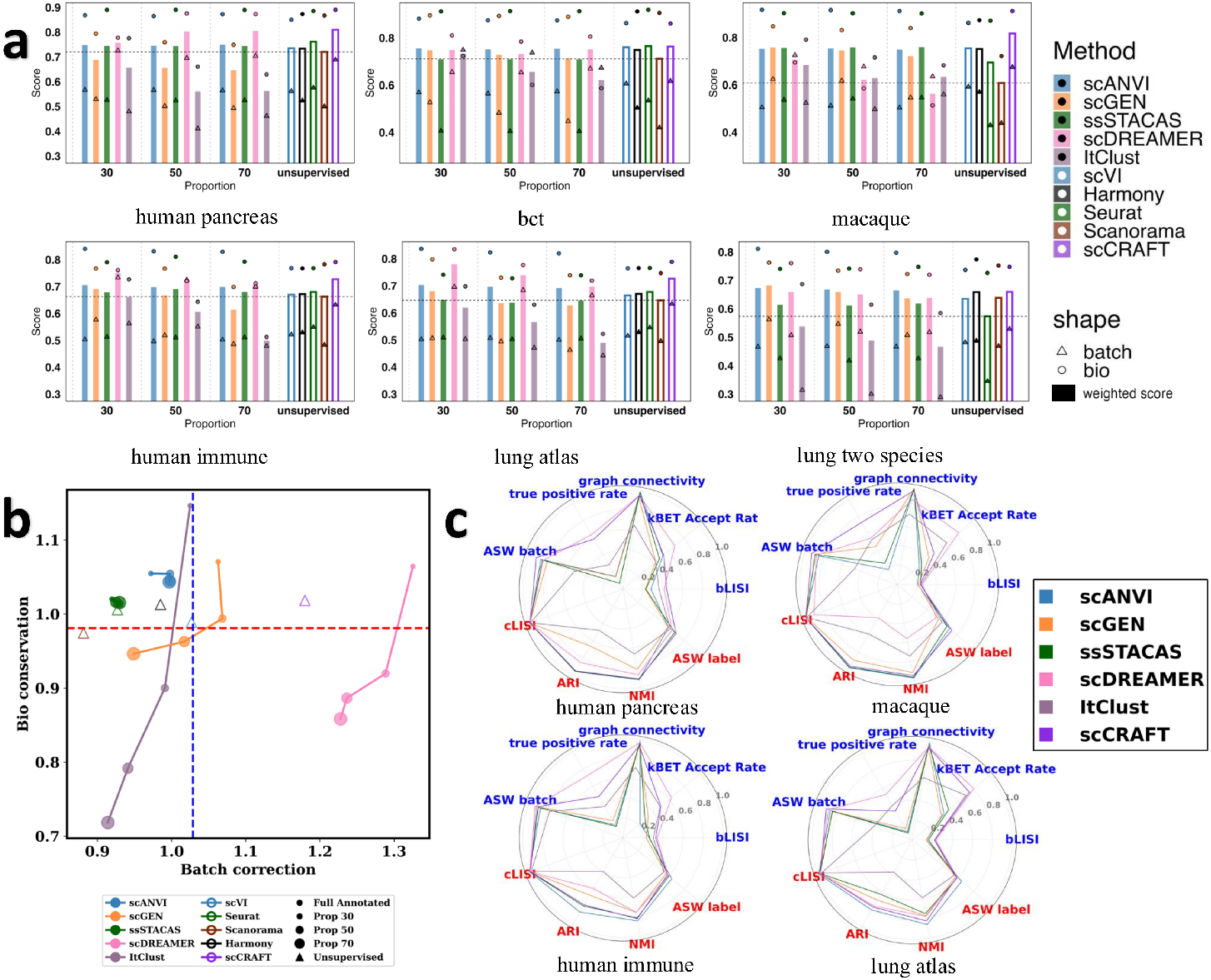
Partial Label Scenario I: Randomly Missing Labels. (**a**) Bar plots showing the performance of all methods across six datasets under this setting. Each bar represents the overall weighted score of a method; triangles and circles indicate the batch correction and biological conservation scores, respectively. The vertical dashed lines divide methods into four groups, namely, 30%, 50%, 70% and unsupervised approaches. The five unsupervised methods are shown on the right, represented by unfilled bars. (**b**) Scatter plot of the scaled batch correction score against the bio-conservation score for each method under the setting for different proportions, averaged across six datasets. The scaled score for each dataset and missing proportion is calculated as the ratio of overall bio-conservation/batch-mixing metric for a given method with respect to the corresponding mean using five unsupervised methods. The detailed scaling procedure can be found in Methods Section 2.2. Scaled scores for unsupervised methods are also included using unfilled triangles. Different colors indicate the methods and the size of dot shapes represent the missing proportions. The horizontal red dashed line represents the average bio conservation score across all methods (both supervised and unsupervised methods), while the vertical blue dotted line represented the average batch correction score. (**c**) Radar plots showing the performance of all methods on individual metrics for the *human pancreas, macaque, human immune*, and *lung atlas* datasets, averaged over all the three proportions for semi-supervised methods. Metrics include biological conservation (red) and batch correction (blue). As scCRAFT achieved the highest overall performance among unsupervised methods, only its scores are shown for clarity; radar plots for the remaining methods are provided in the Supplementary Note 3.

To compare each method’s overall performance across datasets while accounting for datasets difficulty, we scaled the metric scores relative to the performance of unsupervised methods for each dataset and then averaged the scaled scores across datasets for each proportion of missing labels, including performance under the oracle setting as a reference (Fig. 3b). When focusing on biological signal preservation, scANVI ranked first across all levels of label missingness, outperforming the second-best method by 2.48% at 70% missingness after scaling. In contrast, for batch-effect removal, scDREAMER outperformed all other semi-supervised and unsupervised methods, exceeding the second-best method, scCRAFT, by 4.08% at 70% missingness. scCRAFT maintained a balance between the two objectives, with both scores remaining above average. Across all semi-supervised methods, performance declined in both bio-conservation and batch correction as label missingness increased, though the extent varied. Consistent with the unweighted overall performance trends, ssSTACAS and scANVI remained stable, while ItClust showed the steepest decline especially in bio-conservation. From full annotation to 30% missing labels, ItClust’s overall performance dropped by 15.09%; from 30% to 70% missingness, it declined by an additional 15.79%, including a 25.28% drop in bio-conservation and 8.42% in batch correction highlighting its strong dependence on label availability.

From a closer view, we chose four datasets, human pancreas, macaque, human immune and lung atlas datasets as example, covering the relative easiest to hardest datasets, to comprehensively represent the performance of these methods under different scenario. Focusing on methods’ performance on each metric (Fig. 3c), we find that scDREAMER outperforms other methods in true positive rate and kBET Accept rate, surpassing the second best method scCRAFT by on average 6.84% and 23.22%, respectively across the four datasets, which contributes to its superior performance in the batch integration task. Semi-supervised methods did not show significant improvement over scCRAFT when it comes to other metrics.

In Scenario II, we assign incorrect labels to 30%, 50%, and 70% of randomly chosen cells by other cell types in a given dataset. While we observed similar trends as in Scenario I with randomly missing labels, randomly wrong labels are generally more challenging for semi-supervised integration compared to missing labels (See Supplementary Note 4 and Fig. S6). For example, while ssSTACAS and scANVI remained two of the most robust semi-supervised methods, their performance gain over their respective unsupervised backbones (Seurat RPCA and scVI) was less significant than in the randomly missing setting, with their bio-conservation scores no longer consistently surpassing them. Moreover, under the extreme 70% wrong label condition, scDREAMER’s performance on the macaque dataset drops sharply, resulting in an overall score (averaged across all datasets) 15.84% lower than scANVI’s. This places scDREAMER second to last among semi-supervised methods, with scANVI remaining the top performer. In addition, scCRAFT shows better overall performance to semi-supervised methods in most datasets, surpassing scANVI and ssSTACAS by an average 5.05% and 8.20% with 70% of missingness respectively.

### 2.4 Scenario III: Missing and Mixing at Edge

In Scenario III, we mix labels between transcriptionally closest cell types rather than randomizing them. Concretely, in this missing and mixing at edge scenario, we first conduct unsupervised integration with Harmony, then, we examine the 30 nearest neighbors for each cell and compute the proportion of neighbors that share its true cell type. If this proportion falls below a threshold *γ ∈* [0, 1], the cell’s label is reassigned to a different type sampled from its neighbors’ label distribution. If the resampled label still matches the cell’s true type, we instead mark the label as “unknown.” Thus, a higher *γ* results in more label changes due to more cells being selected, and once selected, the cell’s label is changed to either its neighboring cell type or “unknown”, making the scenario more difficult. We set the proportion *γ* = 30%, 50%, 70%. This approach better reflects real-world misannotations, where prediction methods tend to assign cells to transcriptionally similar rather than arbitrary types [15].

Given the same proportion threshold, the actual percentage of reassigned cells varies across datasets, resulting in 1.08% (0.00%), 2.66% (0.88%), 3.91% (1.02%), 12.84% (4.61%), 13.63% (5.15%), and 24.96% (8.84%) reassignments for the BCT, macaque, human pancreas, lung two species, human immune, and lung atlas datasets respectively at *γ* = 70%, with the proportion of missing labels reported in the parenthesis. Since human pancreas, bct and macaque have very small percents of reassignments, we focus on human immune, lung atlas and lung two species in Scenario III. The reassigned proportion in each datasets at *γ* = 30%, 50%, as well as the visualization of Harmony integration results is present in Supplementary Note 5.

On human immune datasets, semi-supervised methods yield performance that are comparable or marginally improved relative to their unsupervised backbones. The comparison between scDREAMER and scCRAFT shows nearly equivalent batch correction score, but a 1.34% advantage for scDREAMER in the bio-conservation score when the reassignment percentage is 13.63% (*γ* = 70%). The performance of scANVI and ssSTACAS was comparable with their respective unsupervised backbones, scVI and Seurat RPCA. scGEN, however, provided a more substantial 2.64% increase in overall performance compared to scVI. This general trend held in the lung two-species datasets, although in this setting, scGEN and scDREAMER both outperformed scCRAFT, by 4.09% and 0.59% (Fig. 4a and c).

**Fig. 4.**
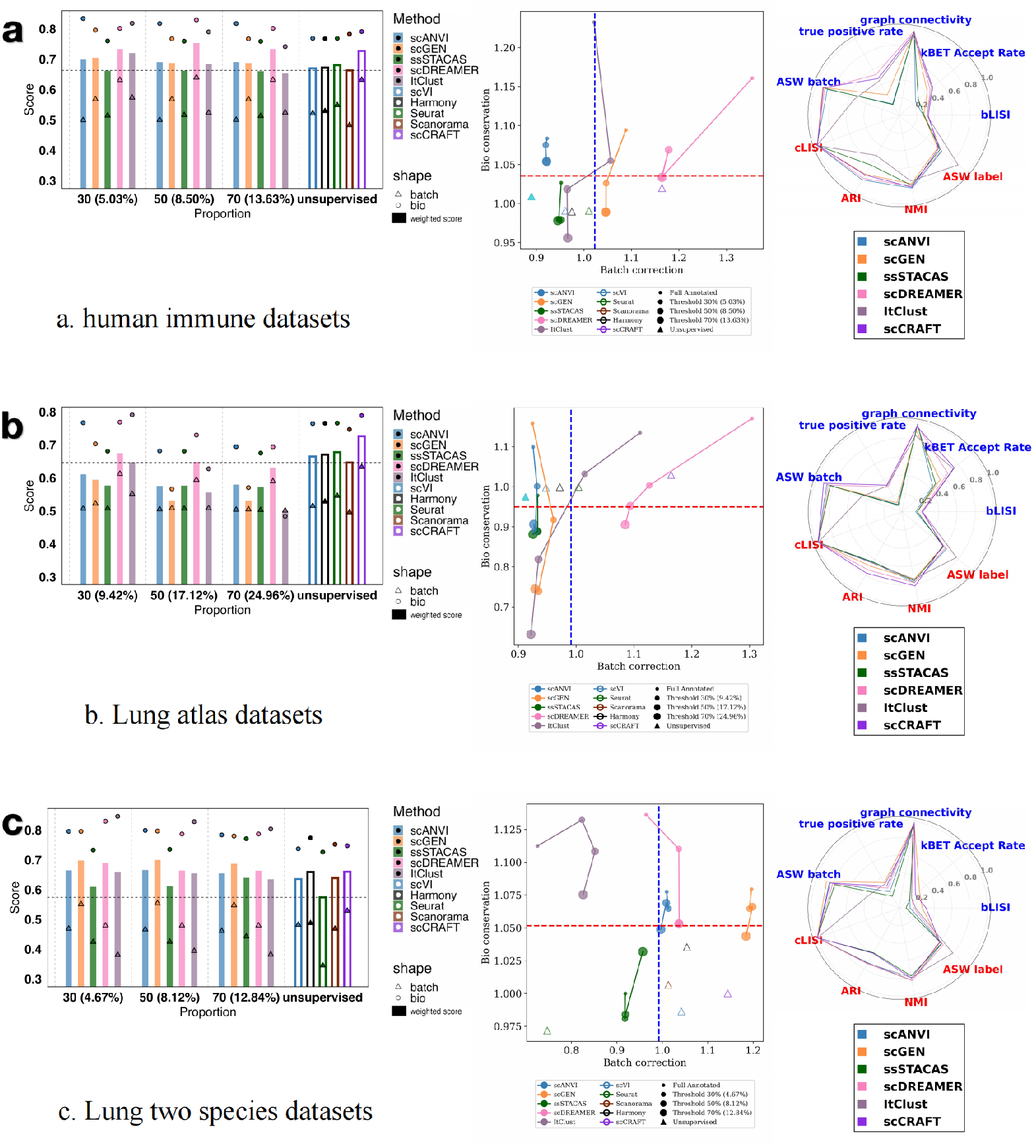
Scenario III: Missing and Mixing at Edge. Plots of semi-supervised and unsupervised methods across three selected datasets under varying levels of label reassignment in Scenario III. Results are shown for three datasets: **(a) *human immune*, (b) *lung atlas***, and **(c) *lung two species***. Each row corresponds to a single dataset and presents a bar plot (left), a scatter plot of scaled scores (middle), and a radar plot of individual metrics (right). **Left Panels (Bar Plots):** The overall weighted score is shown for semi-supervised methods (solid bars) at different label reassignment proportions (controlled by threshold *γ*) and for unsupervised methods (empty bars). Overlaid circles and triangles indicate the biological conservation and batch correction scores, respectively. **Middle Panels (Scatter Plots):** The trade-off between the scaled batch correction score (x-axis) and bio-conservation score (y-axis) is visualized. Methods are color-coded, and the size of the data points corresponds to the reassignment percentage. Solid circles represent semi-supervised results at different *γ* thresholds; empty triangles represent unsupervised methods. The horizontal (red) and vertical (blue) dashed lines mark the mean bio-conservation and batch correction scores, respectively. The scaling procedure is detailed in the Methods Section 2.2. **Right Panels (Radar Plots):** Performance is broken down across individual biological conservation (red) and batch correction (blue) metrics. For semi-supervised methods, scores are averaged across all proportions. For clarity, scCRAFT is shown as a representative high-performing unsupervised method. (Complete radar plots for all methods are provided in Supplementary Note 3).

In contrast, within the lung atlas datasets, which exhibit the highest label reassignment percentage for each *γ* threshold, a substantial performance degradation was observed across all semi-supervised methods. Even when the reassignment rate is below 30%, these methods began to underperform their corresponding unsupervised backbones. This finding suggests that existing semi-supervised approaches may possess a lower tolerance to imperfect labels than previously assumed.

Specifically, at a *γ* value of 70% (reassignment rate = 24.96%), the overall scores for scANVI and scGEN fell below by 7.55% and 22.12% respectively, relative to scVI, while the ssSTACAS score was 11.81% lower than that of Seurat RPCA. Under the same conditions, scDREAMER’s score was 11.40% lower than scCRAFT’s, yet its performance remained comparable to or higher than other unsupervised methods. The scatter plots reveal that all semi-supervised methods exhibit a significant drop in their bio-conservation scores as the reassignment proportion increases. For example, the 10 bio-conservation score for scANVI, a relatively stable method in the previous section, dropped by 2.48% at a 9.42% reassignment rate (*γ* = 30%) and by 2.86% at a 24.96% reassignment rate (*γ* = 70%), relative to the fully annotated baseline. Although ssSTACAS’s performance was stable across different thresholds, its bio-conservation score was consistently inferior to its counterpart, Seurat RPCA. The low bio-conservation scores from semi-supervised methods may be partially attributed to their weaker performance on the ASW label and ARI metrics. Notably, while scDREAMER’s batch correction score also declined, it remained superior to unsupervised methods other than scCRAFT, reaffirming its advantage in this task, possibly due to its high true positive rate and kBET acceptance rate (Fig. 4b).

### 2.5 Scenario IV: Integration with Partially Annotated Batches

Scenario IV reflects a realistic use case in which users have a few well-annotated batches and aim to integrate them with newly acquired, unannotated data. We evaluate whether annotations from the current batches can improve integration performance under various semi-supervised methods. Specifically, we randomly select 30%, 50%, or 70% of the batches to lack cell-type labels (using consistent random seeds across methods and datasets for fair comparison), setting their labels to “unknown”, mimicking incoming unannotated data. Other batches still include the true cell-type labels. For datasets with limited batch numbers such as lung two species (2 batches) and bct (3 batches), we adapted the setting design, and to remain consistency, we move the results for these two datasets to Supplementary Note 6 and exclude these two datasets in the discussion below.

In this setting, scANVI and ssSTACAS are the top-performing semi-supervised methods in this setting, with nearly identical overall scores. While scANVI maintains a slight 0.23% average advantage over ssSTACAS with 70% unannotated batches, both methods significantly outperform other semi-supervised approaches. Notably, ssSTA-CAS surpasses the third-ranked method, scGEN, by 19.57%. Furthermore, scANVI and ssSTACAS only lag behind the top-performing method, scCRAFT, by 8.55% and 8.79% respectively. They outperform their unsupervised counterparts (scVI and Seurat RPCA) by 0.51% and 0.60% respectively under this condition. However, overall performance for semi-supervised methods did not observe significant improvement over unsupervised methods (Fig. 5a).

**Fig. 5.**
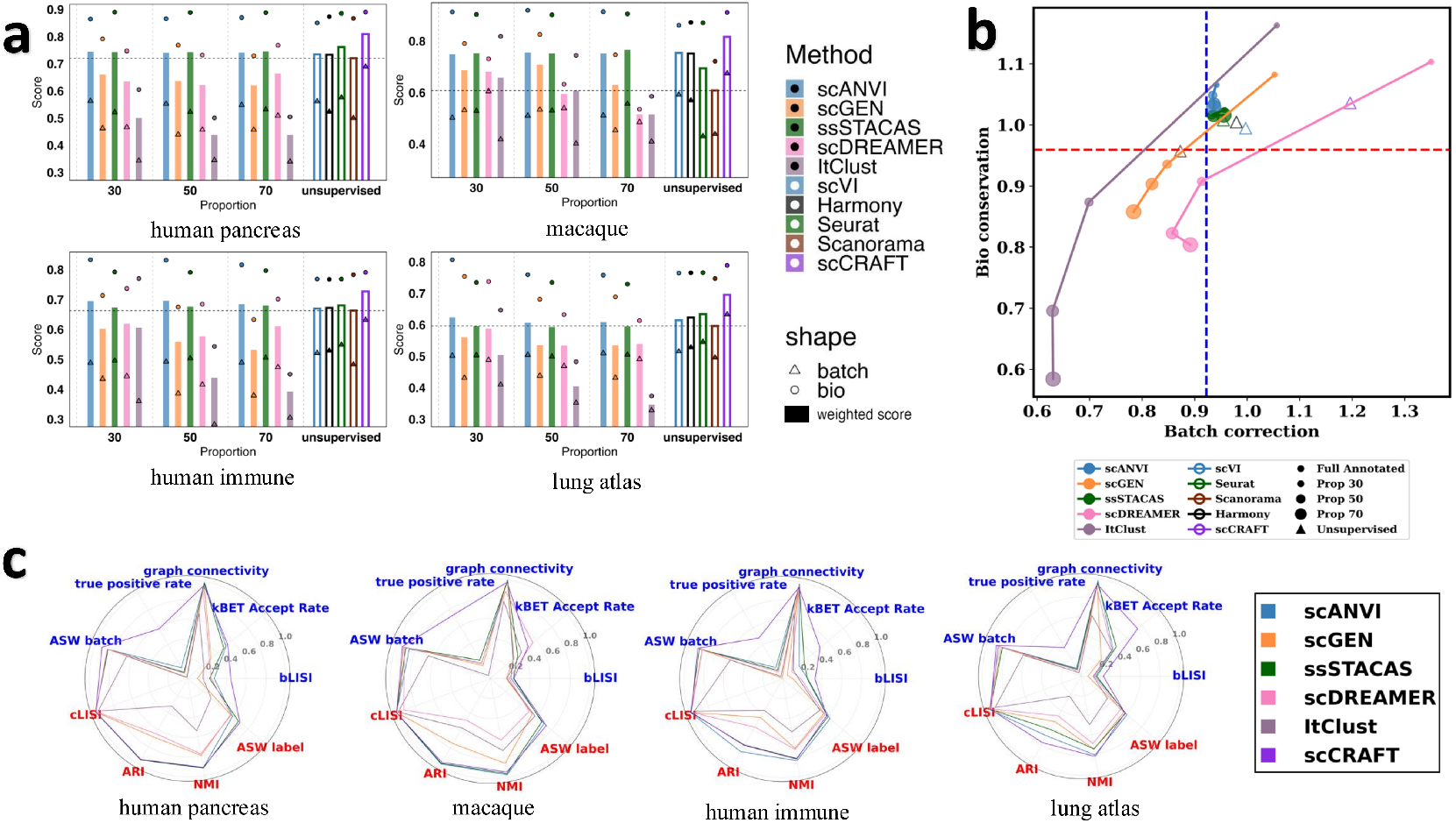
Scenario IV: Partially Annotated Batches. (a) Bar plots of all the methods’ performance in four datasets under this setting. The bar indicates the overall weighted score of each method, while the triangle and circle represents the batch correction scores and bio-conservation scores respectively. The vertical dashed lines separate the bars into four groups, namely, 30%, 50%, 70% and unsupervised. The five unsupervised counterparts are presented on the right, depicted with empty (unfilled) bars. (**b**) Scatter plot of the scaled batch correction score against the scaled bio-conservation score for each method under the partially annotated batches setting for different proportions, averaging across four datasets. The scaled score for each dataset and missing proportion is calculated as the ratio of overall bio-conservation/batch-mixing metric for a given method with respect to the corresponding mean using five unsupervised methods. The detailed scaling procedure can be found in Methods Section 2.2. Scaled scores for unsupervised methods are also included using unfilled triangles. Different colors indicate the methods and the size of dot shapes represent the missing batch proportions. The horizontal red dashed line represents the average bio conservation score across all methods (both supervised and unsupervised methods), while the vertical blue dotted line represented the average batch correction score. (**c**) Radar plots showing the performance of all methods on individual metrics for the *human pancreas, macaque, human immune*, and *lung atlas* datasets, averaged over all the three proportions for semi-supervised methods. Metrics include biological conservation (red) and batch correction (blue). As scCRAFT achieved the highest overall performance among unsupervised methods, only its scores are shown for clarity; radar plots for the remaining methods are provided in the Supplementary Note 3.

Only scANVI and ssSTACAS showed above-average bio-conservation and batch-mixing scores with 30% of unannotated batches (Fig. 5b). Similar as Scenarios I-III, semi-supervised methods showed a drastic drop in bio-conservation performance as the proportion of unannotated batches increases. For example, the bio-conservation score of ItClust, the most label-sensitive approach, falls by 49.57% when the rate of unannotated batches rises from 30% to 70%. Based on the scaled bio-conservation metric scores at 70% of missing batches annotation, scANVI is the leading semi-supervised method, outperforming ssSTACAS by 1.26% and its unsupervised counterpart, scVI, by 3.44%, while lagging just 0.68% behind the best performer, scCRAFT. Regarding batch-effect removal performance, semi-supervised methods generally performed worse than unsupervised ones even with merely 30% of unannotated batches, with only scANVI and ssSTACAS achieving results comparable to some unsupervised methods, but still much worse than the best unsupervised method, scCRAFT. For example, scANVI was surpassed by scCRAFT by 28.02% across all datasets at 30% when examining the scaled batch correction scores. Consistent with the evaluations before, even though scANVI and ssSTACAS were most robust methods in Scenario IV, they did not improve over leading unsupervised approaches, with scDREAMER, ItClust and scGEN’s performance deteriorating rapidly when the proportion of unannotated batches increasing and becoming worse than unsupervised methods with a moderate 30% of missing batches.

When examining individual metrics, we found that scANVI and ssSTACAS outperformed other semi-supervised methods on three of the four biological conservation metrics—ARI, ASW (label), and NMI—and were comparable to or better than the top-performing unsupervised methods on these metrics (Fig. 5c). This explains their strong performance in the overall biological conservation score. However, semi-supervised methods did not show improvement over unsupervised methods on other metrics. Notably, scCRAFT demonstrated a substantial advantage in true positive rate, which largely accounts for its superior overall performance.

### 2.6 Scenario V: Integration with Auto-annotated Labels

As highlighted in the introduction, auto-annotated labels are readily accessible and commonly utilized as auxiliary labels for semi-supervised integration tasks [15]. Therefore, in Scenario V, we benchmark the ability of semi-supervised methods to leverage auto-annotated labels. We consider three widely used auto-annotation tools— SingleR [44], CellAssign [30], and Azimuth [29]—chosen for their popularity and practical usability [17]. CellAssign requires a marker gene matrix, whereas SingleR and Azimuth require reference datasets. Due to limitations in available references, the macaque and BCT datasets are excluded from this setting, as no suitable reference datasets could be identified for them. The predicted cell types from auto-annotation tools vary depending on the reference datasets used and often differ from the original annotations. Cell types not represented in the reference are typically labeled as “other” by CellAssign or “unknown” by SingleR. Below, we summarize the number of cell types predicted by each method across datasets (See Table 1).

**Table 1.** Number of Cell Types Predicted by Auto-Annotation Tools

| Dataset | Original | Azimuth | CellAssign | SingleR |
| --- | --- | --- | --- | --- |
| Human Pancreas | 14 | 13 | 11 | 26 |
| Human Immune | 16 | 29 | 5 | 29 |
| Lung Atlas | 17 | 20 | 8 | 36 |
| Lung Two Species | 17 | 23 | 7 | 37 |

Azimuth predicted labels appear to provide the best annotation quality relative to the original labels—yielding the highest overall scores across all datasets for label-sensitive methods, such as scDREAMER, ItClust, and scGEN. Among semi-supervised approaches, scDREAMER achieved the highest overall weighted score with Azimuth annotations (Fig. 6a). Moreover, scDREAMER performed the best using Azimuth predicted labels among all three auto-annotation labels, achieving average improvements of 12.43% in scaled bio conservation score and 9.46% in scaled batch correction score across all datasets (Fig. 6b).

**Fig. 6.**
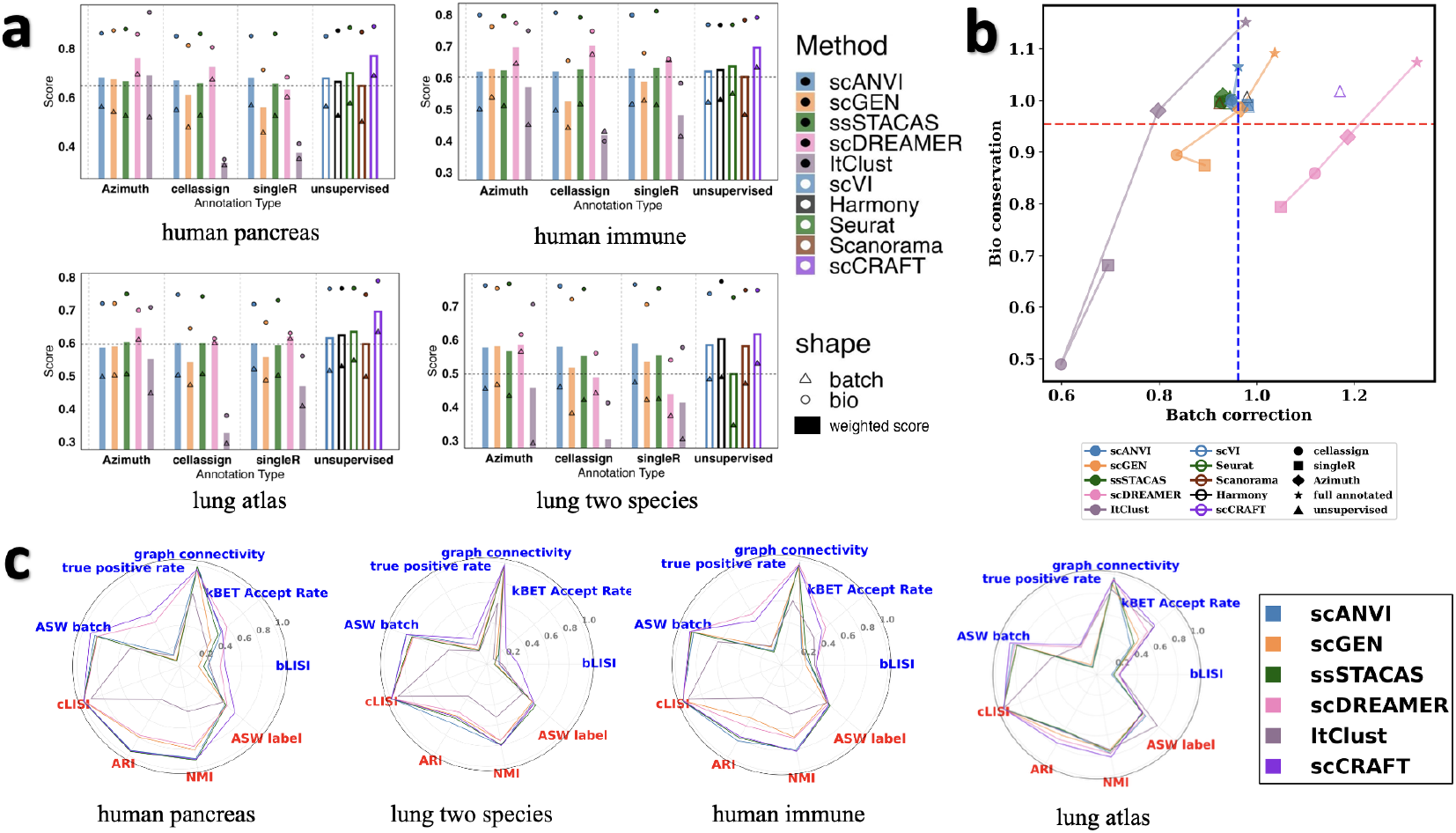
Scenario V: Integration with Auto-annotated Labels. (a) Bar plots showing the performance of all methods across four datasets under this setting. Each bar represents the overall weighted score of a method; triangles and circles indicate the batch correction and biological conservation scores, respectively. The vertical dashed lines divide methods into four groups: those using Azimuth, CellAssign, SingleR, and unsupervised approaches. The five unsupervised methods are shown on the right, represented by unfilled bars. (**b**) Scatter plot of scaled batch correction scores versus biological conservation scores for each method, averaged across the four applicable datasets. Different colors indicate methods, and point shapes represent the origin of the labels. The scaled score for each dataset and auto-annotated labels is calculated as the ratio of overall bio-conservation/batch-mixing metric for a given method with respect to the corresponding mean using five unsupervised methods. The detailed scaling procedure can be found in Methods Section 2.2. The horizontal red dashed line marks the average biological conservation score across all methods, while the vertical blue dotted line marks the average batch correction score. (**c**) Radar plots showing the performance of all methods on individual metrics for the *human pancreas, lung two species, human immune*, and *lung atlas* datasets, averaged over all annotation types for semi-supervised methods. Metrics include biological conservation (red) and batch correction (blue). As scCRAFT achieved the highest overall performance among unsupervised methods, only its scores are shown for clarity; radar plots for the remaining methods are provided in the Supplementary Note 3.

scDREAMER also performed favorably with CellAssign annotations, achieving the highest weighted score among semi-supervised methods in all datasets except for *lung two species*, while its performance was more mixed with SingleR. Both scANVI and ssSTACAS showed robustness to variation in auto-annotation sources (Fig. 6a). However, semi-supervised methods did not consistently outperform unsupervised alternatives (Fig. 6a). For example, scCRAFT outperformed scDREAMER by an average of 5.46% in overall score, respectively, when using Azimuth annotations. scANVI performed comparably to scVI, whereas ssSTACAS performed slightly worse than Seurat RPCA in all datasets except *lung two species*, where it showed a notable improvement.

The scaled metric scores relative to the performance of unsupervised methods within each dataset revealed that scANVI and ssSTACAS consistently performed at levels comparable to unsupervised approaches, with the exception of being lower than scCRAFT in terms of batch correction (Fig. 6b). scGEN and ItClust showed similar bio-conservation performance to scANVI and ssSTACAS when using Azimuth predicted labels, but their performance declined with other auto-annotation sources.

In terms of batch-effect removal, scCRAFT and scDREAMER remained the top two performers. Notably, scDREAMER using Azimuth annotations achieved a batch correction score comparable to scCRAFT, even surpassing it by 1.35%. However, its performance declined with other annotation sources, falling below scCRAFT in both biological conservation and batch mixing. The radar plots show that scDREAMER achieves the highest true positive rate among semi-supervised methods with average performance using three auto-annotation labels (Fig. 6c). Notably, it ranks first in three datasets—except for the lung two-species dataset. However, it was surpassed by the scCRAFT in all four datasets. One important issue using these auto-annotation methods is that the predicted labels have discrepancy compared with the original ones. To assess its possible effect, we do fuzzy matching to compare the results. We include this in our Supplementary Note 7.

## 3 Conclusion

Our results overturn the common expectation that adding cell-type labels automatically improves single-cell integration. With perfect annotations, scDREAMER delivers the best overall compromise between batch mixing and biological conservation, and ItClust best retains the biological signals for defining cell types. The picture changes once labels are less than perfect. A modest 30% random dropout leaves scDREAMER slightly below scCRAFT and scANVI, and any structured imperfection—missing whole batches or label swaps at lineage boundaries—causes scDREAMER, ItClust, and scGEN to fall below not only the strongest unsupervised tool (scCRAFT) but also routine baselines (scVI, Harmony, Seurat RPCA). The same collapse occurs when labels come from popular auto-annotation pipelines (Azimuth, CellAssign, SingleR). Only scANVI-and, to a lesser extent, ssSTACAS—maintains stable performance across all flawed-label scenarios, yet ssSTACAS seldom surpasses its unsupervised counterpart (Seurat RPCA). scANVI’s overall margin over its unsupervised counterpart scVI is small, and neither method ever surpasses scCRAFT.

Although semi-supervised integration might be a promising strategy to improve integration quality by utilizing additional labeling information, unfortunately, at current stage, scCRAFT remains the most reliable “default” integrator in practice when annotation quality is uncertain. scANVI is worth considering only when a modest bump over its parent method scVI is sufficient, or when the main goal is label transfer rather than maximal integration quality. Across every scenario that involved structured label gaps or errors—batch-wise missingness, boundary mis-annotations, or labels generated by automated pipelines—none of the semi-supervised tools examined (scANVI, scGEN, ssSTACAS, scDREAMER, ItClust) delivered a clear advantage over widely used unsupervised options such as scVI, Harmony, or Seurat RPCA.

## 4 Methods

### 4.1 Datasets and Preprocessing

We benchmarked semi-supervised data integration methods on six datasets (See Table 2). Detailed description of each datasets can be found in Supplementary Note 2.

**Table 2.** Summary of Datasets Used in Integration

| Integration Task | Cell Number | Batches | Test Features |
| --- | --- | --- | --- |
| Pancreas | 16382 | 9 | Technologies, donors, rare cell types |
| Bct (mammary epithelial cells) | 9288 | 3 | Small scale with fewer cell types |
| Macaque | 30302 | 30 | Large number of batches |
| Human immune | 33506 | 10 | Tissues, laboratories, similar cell types |
| Lung atlas | 32472 | 16 | Tissues, laboratories, technologies, heterogeneity in cell composition |
| Lung two species | 20760 | 2 | 2-species integration |

Our scRNA-seq data preprocessing follows the established Scanpy [45] pipeline. Before importing the raw count matrices into Scanpy AnnData objects, we filter out low-quality cells with fewer than 300 detected genes and remove genes detected in fewer than 5 cells to avoid misleading alignments caused by dropout events or low transcriptional activity. We then store the raw counts in the count layer so that models like scVI and scANVI, which require raw count input, remain unaffected by later transformations. Next, each cell’s library size is normalized to 10,000 reads, scaling gene counts by total counts per cell and adjusting them to a common scale, followed by a log transformation using log(1+x) to stabilize variance across the dataset. After these steps, we identify the top 2,000 highly variable genes to capture the dataset’s biological diversity while correcting for batch effects.

This preprocessing pipeline is uniformly applied across our benchmarking methods except for scVI, scANVI, scDREAMER and ItClust. While scVI and scANVI are configured to use the preserved raw counts, scDREAMER and ItClust employ its own self-preprocessing function. scDREAMER’s self-preprocessing function is designed to take an entire h5ad file as input and then using Scanpy to do normalization and log transformation, using the same parameters as we did. It then selects the top 2,000 highly variable genes using the Seurat RPCA method, with batch correction handled via the provided batch key. Beyond these standard steps, scDREAMER further converts the data into either a dense or sparse matrix (depending on the configuration set by sparseIP) and one-hot encodes batch information and cell type labels for downstream model use.

ItClust includes preprocessing steps, that is, filtering of cells/genes, normalization, scaling and selection of highly variables genes, similar to what we did for other methods, but with different parameters. It filters out low-quality cells with fewer than 200 detected genes and remove genes detected in fewer than 10 cells. Besides, the normalization is done to 100,000 reads, and the top 1,000 highly variable genes are selected.

Following preprocessing, we adhere to the standard benchmarking protocols for each method, with detailed procedures outlined in Supplementary Note 1.

### 4.2 Evaluation Metrics

#### 4.2.1 Overview

In our analysis of single-cell data integration methods, we employed a structured approach to evaluate performance, drawing upon established frameworks in recent studies. We focused on two critical aspects: (1) the conservation of biological variance and (2) the correction of batch effects. Following the methodology outlined in contemporary research, we selected global cluster matching (which includes the Adjusted Rand Index (ARI) and normalized mutual information (NMI), silhouette width by cell-type label (ASW label) and cLISI as four key metrics for assessing the conservation of biological variance. These metrics provide insights into how well each method preserves essential biological information and cell identity. For the correction of batch effects, our evaluation incorporated five specific metrics: the average silhouette width across batches (ASW batch), the k-nearest-neighbor batch effect test (kBET Accept Rate), bLISI, k-nearest-neighbor graph connectivity, and true positive rate. These metrics collectively assess the effectiveness of each integration method in harmonizing data across different batches, ensuring that technical variations do not obscure biological signals. Overall accuracy scores were computed by taking the weighted mean of all metrics computed for an integration run, with a 60/40 weighting of biological variance conservation (bio-conservation) to batch effect removal irrespective of the number of metrics computed.

#### 4.2.2 Scaling Procedure For Scatter Plots

In panel (b) of Figure 3, 4, 5, 6, we use the scaled score for bio-conservation and batch correction for scatter plots. The scaled score is calculated through the following steps:

- **Compute Unsupervised Baselines:** For each dataset and score type (bio or batch), calculate the mean score of all unsupervised methods. This establishes a baseline for each dataset and score type.
- **Normalize Raw Scores:** For every unique combination of dataset, labeled data proportion, and method (including unsupervised ones), normalize the method’s raw score by dividing it by the corresponding unsupervised baseline (i.e., the baseline for the same dataset and score type).
- **Average Scaled Scores:** For each combination of method, labeled data proportion level, and score type, compute the average of the normalized scores obtained in Step 2 across all datasets.

#### 4.2.3 ARI

The adjusted Rand index (ARI) measures agreement between two partitions by adjusting the raw Rand index for chance overlap. In our workflow, we compared true cell-type annotations against Louvain clusters derived from the integrated scRNA-seq data. By accounting for random label assignments, the ARI yields 0 for clusterings no better than chance and 1 for perfect concordance. We computed ARI using the scib package.

#### 4.2.4 NMI

The normalized mutual information (NMI) quantifies the similarity between two clusterings by measuring the amount of shared information, normalized to account for differences in cluster sizes. We used NMI to evaluate how well Louvain clusters derived from the integrated data align with known cell-type annotations. An NMI of 0 indicates no mutual information (random agreement), while a value of 1 reflects perfect overlap. All NMI scores were calculated using the scib package.

#### 4.2.5 Cell Type Average Silhouette Width (ASW label)

The average silhouette width (ASW) quantifies how well-defined clusters are by comparing the average within-cluster distance of a cell to its average distance from the nearest neighboring cluster. Scores range from -1 to 1, with higher values indicating more distinct and cohesive clusters.

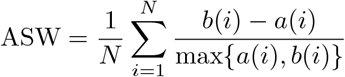

To evaluate biological conservation, we computed ASW based on known cell-type labels (ASW label) using the integrated embedding. An ASW near 1 suggests strong preservation of cell-type structure, whereas values near 0 or negative reflect overlapping or misclassified clusters. To facilitate comparison across methods, we linearly scaled the ASW label scores to a 0–1 range using the formula:

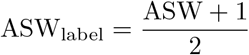

We compute this metric using scib package.

#### 4.2.6 Cell Type Local Inverse Simpson’s Index (cLISI)

The cell-type Local Inverse Simpson’s Index (cLISI) quantifies the diversity of cell types within local neighborhoods to assess biological variance preservation in integrated data. It is computed by applying the inverse Simpson’s Index to the distribution of cell-type labels among each cell’s nearest neighbors, capturing the effective number of distinct cell types present locally. Letting *B* represents the number of unique cell types, and *p*_*b*_ denote the probability of observing cell type *b* in a neighborhood, cLISI is defined as:

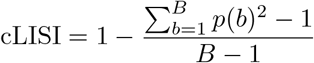

The final cLISI score is averaged across all cells and then min-max scaled to the range [0, 1], where higher values reflect better conservation of biological diversity. This metric is particularly useful for detecting whether cell-type structure is preserved locally after integration.

#### 4.2.7 Average Silhouette Width for Batches (ASW batch)

The batch average silhouette width (ASW batch) assesses how well batches are mixed within each biological cell type after integration. It is based on the absolute silhouette width computed on batch labels, where a value close to 0 indicates good batch mixing, and larger values reflect residual batch effects. To ensure that higher scores correspond to better integration, the silhouette values are subtracted from 1. Specifically, the batchASW for each cell type is computed by averaging 1 *−* |*s*(*i*)|, where *s*(*i*) is the silhouette score of cell *i* with respect to batch labels. These cell-type–specific scores are then averaged across all cell types to obtain the final batch ASW: batch ASW_*j*_

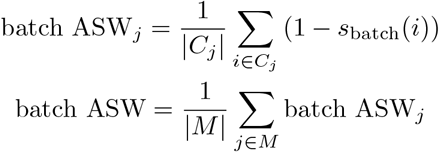

where *C*_*j*_ is the set of cells of cell type *j*, and *M* is the set of all unique cell types. A final batch ASW score close to 1 indicates ideal batch mixing, while a score near 0 implies poor integration. We compute this metric using scib package.

#### 4.2.8 k-Nearest Neighbor Batch Effect Test (kBET)

The k-nearest neighbor Batch Effect Test (kBET) evaluates local batch label composition against the global batch distribution to assess batch mixing. The test is applied iteratively to randomly sampled subsets of cells, with a chi-squared test determining whether local neighborhoods deviate significantly from expected label proportions. For each embedding or corrected feature space, k-nearest neighbor graphs are constructed with a fixed neighborhood size to ensure consistency across methods. When applied, kBET accounts for technical variation by testing batch-wise and adapting the neighborhood size (typically between 10 and 100) based on local connectivity. For disconnected graphs, the test is limited to connected components; if more than 25% of cells belong to components too small for testing, a default score of 1 is assigned, indicating inadequate correction. The final kBET score is computed as the inverse of the average rejection rate across tested cells, where values near 1 indicate well-mixed batches and values near 0 reflect poor integration. All kBET scores were computed using the kBET R package (v.0.99.6).

#### 4.2.9 Batch Label Identity Score Index (bLISI)

The batch Local Inverse Simpson’s Index (bLISI) assesses batch mixing by quantifying the diversity of batch labels within each cell’s local neighborhood. A higher bLISI score indicates that a cell’s neighbors are drawn evenly from multiple batches, reflecting effective batch correction. Ideally, well-integrated datasets yield bLISI values approaching the total number of batches, suggesting uniform local batch composition. To normalize the score across datasets with varying batch counts, bLISI is scaled using the transformation:

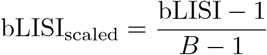

where *B* is the total number of batches. This scaling bounds the score between 0 and 1, where values near 1 indicate optimal batch mixing and values near 0 suggest poor integration.

#### 4.2.10 Graph Connectivity

Graph connectivity evaluates whether cells of the same biological identity remain connected in the integrated k-nearest neighbor (kNN) graph, serving as an indicator of biological conservation. For each cell type, the method constructs a subgraph from the global kNN graph by retaining only cells of that type, and then computes the size of the largest connected component relative to the total number of cells of that type. The final graph connectivity score is obtained by averaging these proportions across all cell types:

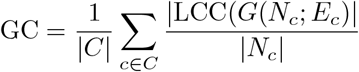

where *C* represents the set of cell identity labels, |LCC()| is the number of nodes in the largest connected component of the graph and *N*_*c*_ is the number of nodes with cell identity |*Nc*|. A score close to 1 indicates that cells of the same type are well-connected post-integration, reflecting strong preservation of biological structure.

#### 4.2.11 Proportion of True Positive Cells (True Positive Rate)

The true positive rate (TPR) evaluates integration quality at the single-cell level by assessing the local consistency of cell type and batch composition. A cell is considered positive if its local neighborhood consists predominantly of cells from the same annotated type. Among these, true positives are those whose local batch distribution also reflects the global batch proportions, indicating proper batch mixing. The TPR is then defined as the fraction of true positives among all positive cells, capturing both biological variance conservation and batch effect correction. Higher TPR values signify that the integration method preserves cell identity while achieving balanced batch mixing in local neighborhoods.

#### 4.2.12 Overall scores and metric aggregation

Metrics were run on the integrated embeddings output by each method. The overall score, *S*_overall,*i*_, for each integration run *i* was calculated by taking the weighted mean of the batch removal score, *S*_batch,*i*_, and the bio-conservation score, *S*_bio,*i*_, following the equation:

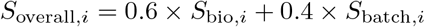

Here we assign higher weights to bio-conservation score since we focus more on semi-supervised methods’ performance on the preservation of biological variations.

Specifically, the partial scores were computed by averaging all raw metrics that belong to each type via:

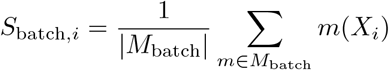

and

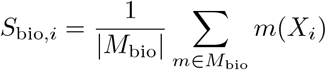

Here, *X*_*i*_ denotes the integration output for run *i*, and *M*_batch_ and *M*_bio_ denote the set of metrics belong to batch removal and bio-conservation scores respectively.

## Acknowledgements

This work is supported by NSF DMS2310836 and NIH 5U01AI167892.

## Declarations

- Data availability: See Supplementary Material, Suppl Section 2
- Code availability: https://github.com/RainySheena/benchmark_semi

