## Supplementary file for "A Benchmark of Semi-Supervised scRNA-seq Integration Methods in Real-World Scenarios"

### Benchmarking of semi-supervised scRNA-seq integration methods in realistic settings

**3 Radar Plots for all Methods in Specific Datasets** **8**

**4 Scenario II: Randomly Wrong Labels** **12**

**5 Scenario III: Harmony Integration Results of Each Datasets Under Different Proportion Thresholds** **14**

**6 Scenario IV: Partially Annotated Batches Results for Lung Two Species and BCT Datasets** **17**

**7 Scenario V: Semi-Supervised Integration Methods' Performance on Auto-annotation Labels After Fuzzy Matching** **17**

### 1 Integration Methods

We ran the 10 selected data integration methods, five unsupervised and five supervised methods, according to default parameterizations obtained from available tutorials and paper methods.

To visually represent architectural similarities, methods are colored according to their backbone structure. Consequently, scVI and scANVI are designated with the same blue color, while Seurat RPCA and ssSTACAS share a green color. An exception is made for scGEN, which, despite being built upon the scVI backbone, is allocated a distinct color to emphasize its unique identity relative to scANVI. Methods not belonging to these families are each assigned a unique color. Further details on the algorithms of each method and how each method was run can be found in the section below.

#### 1.1 scVI

The scVI (single-cell Variational Inference) model integrates the principles of a variational autoencoder with a hierarchical Bayesian framework to analyze single-cell gene expression data. By modeling count data using a negative binomial distribution, scVI captures both biological variability and technical noise, including differences in sequencing depth and batch effects. It learns a low-dimensional latent representation of the data that reflects the underlying biological signal while mitigating unwanted variation.

For implementation, we followed the standard workflow recommended in the scVI-tools documentation, accessible at "https://docs.scvi-tools.org/en/stable/tutorials/notebooks/scrna/harmonization.html". The scVI model was initialized with two hidden layers, a 50-dimensional latent space, and a negative binomial gene likelihood. After training the model, we extracted the latent embedding and stored it, which was subsequently used for downstream analyses including nearest-neighbor graph construction and UMAP visualization.

#### 1.2 Harmony

The Harmony algorithm begins by projecting all datasets into a shared PCA space, incorporating batch variables to account for technical variability across samples. It then iteratively alternates between two main steps until convergence. The first step applies a maximum diversity clustering strategy, which prevents overcorrection by encouraging separation between similar cell types from different batches. The second step fits a linear mixture model to adjust for batch effects and update the embedding. This iterative process yields a corrected low-dimensional representation that preserves biological variation while minimizing batch-driven differences.

In our study, we applied Harmony (version 1.0) following the procedures outlined in its official Python tutorial. We used 50 principal components as input and corrected for batch effects using the specified batch variable. The resulting Harmony embedding was then used for downstream analyses including nearest-neighbor graph construction and UMAP visualization.

#### 1.3 Seurat RPCA

The Seurat RPCA integration framework utilizes a reciprocal PCA (RPCA)-based approach to align datasets from different batches by identifying shared cell states across samples. This method begins by individually normalizing each batch using SCTransform, a variance-stabilizing transformation that effectively handles differences in sequencing depth and technical noise. After performing PCA on the normalized data, Seurat RPCA identifies mutual nearest neighbors across batches in the shared PCA space and uses these anchors to integrate the datasets.

In our study, we followed the standard Seurat RPCA integration pipeline, using SCTransform for normalization, 50 principal components for dimensionality reduction, and RPCA as the integration method. The resulting integrated embedding, stored in the "integrated.rpca" reduction slot, was then used to construct a nearest-neighbor graph and generate UMAP visualizations.

#### 1.4 Scanorama

The Scanorama algorithm builds upon the mutual nearest neighbors (MNN) concept, adopting a strategy akin to panoramic image stitching to integrate single-cell datasets. It begins by performing a k-nearest neighbor search to identify analogous cells across batches, then refines these links to a core set of mutual nearest neighbors, which serve as anchors for alignment. Scanorama subsequently projects all datasets into a shared low-dimensional space by aligning them within a unified hyperplane, effectively correcting for batch effects while preserving biological structure.

For implementation, we utilized the "scanorama.correct_scanpy" function, first segmenting the dataset by batch. We enabled the "return_dimred=True" option to obtain both a joint embedding and a batch-corrected expression matrix. The corrected data was then reordered to preserve the original cell order and used for downstream analyses.

#### 1.5 scCRAFT

The scCRAFT framework integrates single-cell RNA-seq data by combining three components to produce robust, batch-invariant embeddings while preserving critical biological structure. It first employs a customized variational autoencoder (VAE) to reconstruct the observed count matrix from a latent embedding space, capturing meaningful variations in the input data. To reduce batch-induced discrepancies, scCRAFT introduces a multi-domain GAN loss, which encourages mixing across batches through domain adaptation. However, to avoid overcorrection—especially in datasets with rare or heterogeneous cell types—the model incorporates a dual-resolution triplet loss. This component leverages the within-batch topological structure by using low- and high-resolution clustering to select informative triplets for contrastive learning.

In our implementation, we first performed multi-resolution clustering on the input data and then trained the integration model using a latent space of 256 dimensions and a domain adaptation coefficient of 0.2. The resulting scCRAFT embedding was used to construct the nearest-neighbor graph and generate UMAP visualizations for downstream analysis.

#### 1.6 scANVI

The scANVI model extends scVI by incorporating semi-supervised learning to jointly perform batch correction and cell type classification. Built on the same variational autoencoder (VAE) architecture as scVI, scANVI introduces an additional classification head that leverages available cell type labels while simultaneously learning from unlabeled data. This approach allows the model to infer biologically meaningful latent representations that are both batch-invariant and informative for cell identity.

In our implementation, we first utilized the trained scVI model before to initialize scANVI, which was further trained using cell type annotations. We trained the model for 20 epochs, sampling 100 cells per labeled class. The resulting scANVI embedding was used for downstream neighbor graph construction and UMAP visualization, while predicted cell types were stored for evaluation.

#### 1.7 scGEN

The scGEN model is a generative deep learning framework designed to address batch effects in single-cell transcriptomic data through a variational autoencoder (VAE) combined with adversarial training. By conditioning on both batch labels and cell type information, scGEN learns a batch-invariant latent representation that enables effective integration across datasets while preserving cell identity.

In our implementation, we initialized the model using annotated training data with both batch and cell type labels. The model was trained for up to 100 epochs with a batch size of 32, incorporating early stopping with a patience of 25 epochs to prevent overfitting. After training, we applied the "batch_removal" function to generate a corrected dataset. The resulting batch-corrected embeddings, stored in the "latent" space, were extracted and used for downstream analysis and visualization.

#### 1.8 ssSTACAS

ssSTACAS is a semi-supervised batch correction method for single-cell RNA-seq data that leverages prior cell type information to guide integration while preserving meaningful biological variability. By incorporating partial or imprecise cell type labels, ssSTACAS enhances the accuracy of anchor identification across datasets and mitigates the risk of overcorrection—an issue particularly prevalent in datasets with rare or non-overlapping cell populations.

In our implementation, we applied ssSTACAS using 2,000 anchor features and 50 principal components, specifying the cell type annotations with a confidence threshold of 0.5. After integration, the data was scaled and PCA was performed to derive the final low-dimensional embedding. The embedding was then stored to match the original cell order for downstream analysis.

#### 1.9 scDREAMER

scDREAMER has unsupervised and supervised version, and here we use supervised version "scDREAMER-Sup". The scDREAMER framework is a deep generative model designed for supervised integration of single-cell RNA-seq data, combining autoencoders with adversarial training to correct batch effects while preserving meaningful biological variation. scDREAMER-Sup leverages cell type labels during training to guide the learning of a biologically informative, batch-invariant latent space.

In our implementation, we applied scDREAMER-Sup to multiple cell type–annotated datasets. For relatively small datasets, we used a learning rate of 0.0002 for the autoencoder and 0.0007 for the discriminator. For larger datasets with more than 0.5 million cells, we adjusted the rates to 0.0001 for the autoencoder and 0.00001 for the discriminator, following the recommended configuration. The model was trained for 240 epochs within a TensorFlow session configured with GPU memory growth control. After training, the embedding was automatically saved and subsequently used for downstream analysis.

##### 1.10 ItClust

ItClust is an iterative transfer learning algorithm designed for clustering and cell type classification in single-cell RNA-seq data. It first trains a neural network on a well-annotated source dataset to learn gene expression signatures and then transfers this knowledge to a target dataset by initializing a second network with the learned parameters. The target network is subsequently fine-tuned in an unsupervised manner to adapt to the unique characteristics of the target data, allowing it to capture target-specific gene expression patterns. This iterative process enables accurate cell clustering even in the presence of strong batch effects or previously unseen cell types.

In our implementation, we followed the ItClust tutorial, available at https://github.com/jianhuupenn/ItClust/blob/master/tutorial/tutorial.md. After training, the learned embedding was saved and used for downstream analyses.

#### 2 Datasets Details

##### 2.1 Lung atlas Datasets

The lung atlas utilized in this study originates from the comprehensive research by Vieira Braga et al. and is publicly accessible via the Gene Expression Omnibus (GEO) database under accession code GSE130148 [https://www.ncbi.nlm.nih.gov/geo/query/acc.cgi?acc=GSE130148]. This dataset provides a valuable resource for investigating the cellular landscape of the lung. The detailed methodology for the dataset's compilation, including criteria for sample selection, the diversity of experimental protocols employed, and specifics concerning data retrieval, is meticulously described in the benchmarking study by Luecken et al.

##### 2.2 Human Pancreas Datasets

The human pancreas dataset utilized in this study is a compilation of five publicly available pancreatic islet datasets, totaling 16,382 cells. Data were sourced from the Gene Expression Omnibus (GEO) database under accession codes GSE81076 [https://www.ncbi.nlm.nih.gov/geo/query/acc.cgi?acc=GSE81076], GSE85241 [https://www.ncbi.nlm.nih.gov/geo/query/acc.cgi?acc=GSE85241],

GSE86469 [https://www.ncbi.nlm.nih.gov/geo/query/acc.cgi?acc=GSE86469], GSE84133 [https://www.ncbi.nlm.nih.gov/geo/query/acc.cgi?acc=GSE84133], and GSE81608 [https://www.ncbi.nlm.nih.gov/geo/query/acc.cgi?acc=GSE81608], and from the ArrayExpress database under accession code E-MTAB-5061 [https://www.ebi.ac.uk/biostudies/arrayexpress/studies/E-MTAB-5061]. The detailed compilation process, encompassing dataset selection criteria and the range of experimental protocols employed, is thoroughly documented in the benchmarking study by Luecken et al. This documentation promotes transparency and supports the reproducibility of analyses utilizing this aggregated dataset.

#### 2.3 Human Immune Datasets

The Human Immune dataset utilized herein comprises cells from ten human samples, representing bone marrow and peripheral blood tissues. Bone marrow samples were sourced from the study by Oetjen et al., whereas peripheral blood samples were compiled from multiple sources, including publicly available data from 10x Genomics (PBMC10k) and studies by Freytag et al., Sun et al., and Villani et al. These source datasets are accessible via the Gene Expression Omnibus (GEO) database under accession codes GSE120221 [https://www.ncbi.nlm.nih.gov/geo/query/acc.cgi?acc=GSE120221], GSE107727 [https://www.ncbi.nlm.nih.gov/geo/query/acc.cgi?acc=GSE107727], GSE115189 [https://www.ncbi.nlm.nih.gov/geo/query/acc.cgi?acc=GSE115189], GSE128066 [https://www.ncbi.nlm.nih.gov/geo/query/acc.cgi?acc=GSE128066], and GSE94820 [https://www.ncbi.nlm.nih.gov/geo/query/acc.cgi?acc=GSE94820], and from the 10X Genomics website for the PBMC10k dataset (https://support.10xgenomics.com/single-cell-gene-expression/datasets/3.0.0/pbmc_10k_v3). Comprehensive details on the procurement of these diverse datasets, the experimental protocols employed, and sample selection criteria are provided in the work by Luecken et al. Following quality control measures applied in the present study, the resulting dataset for analysis was refined to 33,506 cells and 12,303 genes, distributed across ten distinct batches.

#### 2.4 Macaque Datasets

The Macaque Retina dataset, generated by Lakkis et al., serves as an important resource for exploring retinal cell compositions in macaques. This dataset was originally deposited in the Gene Expression Omnibus (GEO) database under accession code GSE11848 [https://www.ncbi.nlm.nih.gov/geo/query/acc.cgi?acc=GSE11848]. Subsequently, the dataset was processed by Yu et al. and is accessible via Figshare at "https://figshare.com/articles/dataset/Batch_Alignment_of_single-cell_transcriptomics_data_using_Deep_Metric_Learning/20499630/2".

#### 2.5 Lung Two Species Datasets

The Lung Two Species dataset, designed to facilitate cross-species comparisons, comprises lung cells from both mouse and human, segregated into two distinct batches. This dataset was originally reported by Raredon et al. and is accessible via the Gene Expression Omnibus (GEO) database under accession code GSE133747 [https://www.ncbi.nlm.nih.gov/geo/query/acc.cgi?acc=GSE133747]. Subsequently, this dataset was processed by Yu et al. and is available on Figshare at "https://figshare.com/articles/dataset/Batch_Alignment_of_single-cell_transcriptomics_data_using_Deep_Metric_Learning/20499630/2". Following quality control procedures, the dataset utilized for analysis contains 20,760 cells and 62,781 genes.

##### 2.6 Bct (Mammary Epithelial Cell) Datasets

The Bct dataset comprises mammary epithelial cells collated from studies by Giraddi et al., Pal et al., and Bach et al. Source data from these contributions are accessible via the Gene Expression Omnibus (GEO) database under accession codes GSE111113 [https://www.ncbi.nlm.nih.gov/geo/query/acc.cgi?acc=GSE111113], GSE98131 [https://www.ncbi.nlm.nih.gov/geo/query/acc.cgi?acc=GSE98131], GSE103275 [https://www.ncbi.nlm.nih.gov/geo/query/acc.cgi?acc=GSE103275], and GSE106273 [https://www.ncbi.nlm.nih.gov/geo/query/acc.cgi?acc=GSE106273]. The compiled dataset is organized into three batches—denoted as spk, vis, and wal—each sharing three cell types: basal, lumina_mature, and luminal_progenitor. This aggregated dataset was subsequently processed by Yu et al. and is available on Figshare at basal, lumina_mature, and luminal_progenitor.

#### 3 Radar Plots for all Methods in Specific Datasets

For clarity, in the main text, we only include the best performed unsupervised method – scCRAFT in the radar plots. Here we show the radar plots with all methods present for the given datasets in each setting.

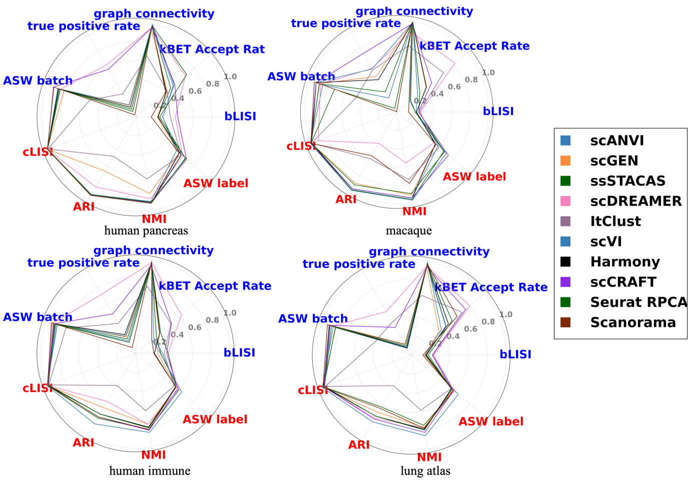

**Fig. S1** Radar Plots for Randomly Missing Setting
Radar plot of all the methods' performance on each metric in human pancreas, macaque, human immune and lung atlas datasets under the randomly missing setting, averaging across all the three proportions, where the metrics include bio-conservation metrics (shown in red) and batch correction metrics (shown in blue).

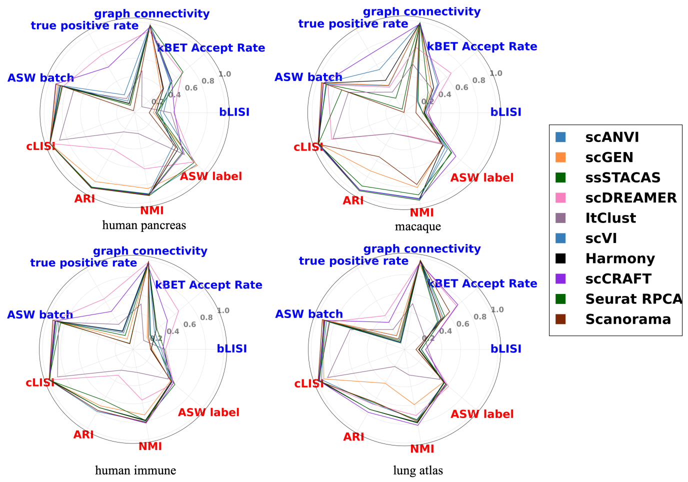

**Fig. S2** Radar Plots for Randomly Wrong Setting
Radar plot of all the methods' performance on each metric in human pancreas, macaque, human immune and lung atlas datasets under the randomly wrong setting, averaging across all the three proportions, where the metrics include bio-conservation metrics (shown in red) and batch correction metrics (shown in blue).

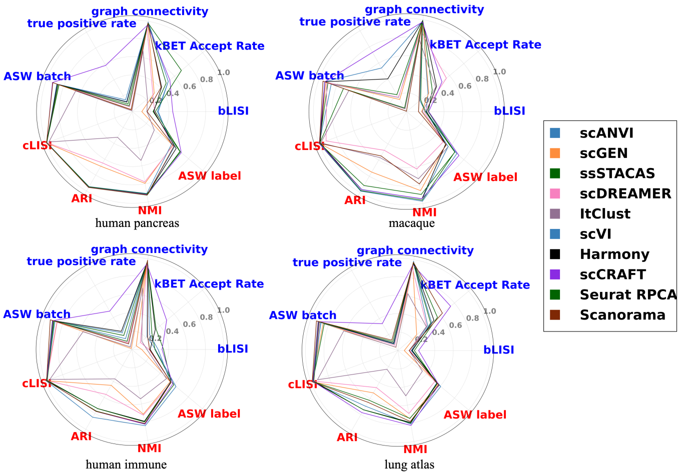

**Fig. S3** Radar Plots for Partially Annotated Batches Setting
Radar plot of all the methods' performance on each metric in human pancreas, macaque, human immune and lung atlas datasets under the partially annotated batches setting, averaging across all the three proportions, where the metrics include bio-conservation metrics (shown in red) and batch correction metrics (shown in blue).

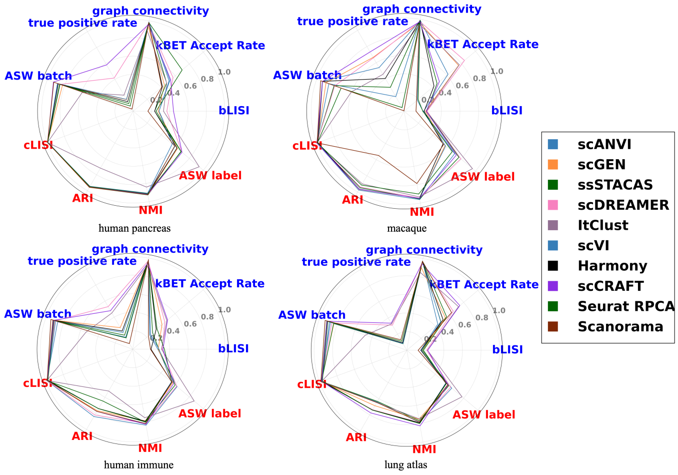

**Fig. S4** Radar Plots for Missing and Mixing at Edge Setting
Radar plot of all the methods' performance on each metric in human pancreas, macaque, human immune and lung atlas datasets under the missing and mixing at edge setting, averaging across all the three proportions, where the metrics include bio-conservation metrics (shown in red) and batch correction metrics (shown in blue).

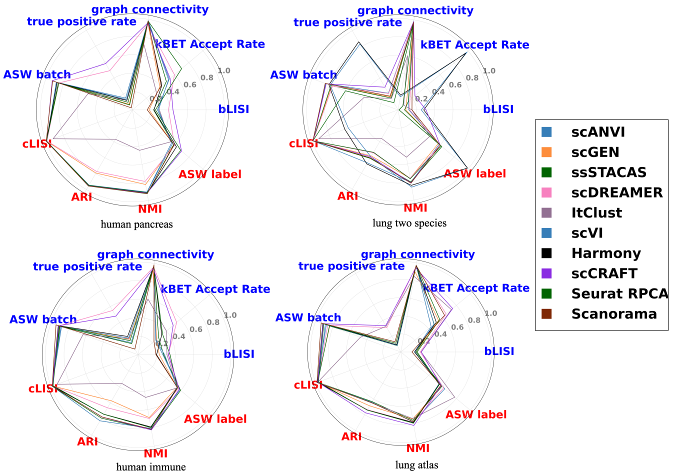

**Fig. S5** Radar Plots for Auto-annotation Labels Setting
Radar plot of all the methods' performance on each metric in human pancreas, lung two species, human immune and lung atlas datasets under the auto-annotation labels setting, averaging across all the three proportions, where the metrics include bio-conservation metrics (shown in red) and batch correction metrics (shown in blue).

### 4 Scenario II: Randomly Wrong Labels

In this benchmark, we evaluated the robustness of semi-supervised integration methods to label noise by randomly assigning incorrect cell-type labels to 30%, 50%, and 70% of the cells in each dataset. This strategy, commonly adopted in benchmarking semi-supervised methods, differs from the missing-label setting: instead of assigning the selected cells an "Unknown" label, we randomly replaced their true labels with incorrect ones. Each incorrect label was chosen uniformly at random from all possible cell types in the dataset, excluding the cell's true label. We applied this perturbation to compare five semi-supervised methods and assess their ability to handle mislabeled annotations.

In this setting, scANVI emerged as the best performer among all the semi-supervised methods considering the overall score in most of the datasets, but its average performance across all datasets at 70% was surpassed by scCRAFT by 5.05%. In general, supervised methods did not observe significant performance improvement over the unsupervised methods (Figure S6a).

To compare the performance of each methods generally across datasets, we scaled the metrics score based on unsupervised methods' performance then average across all datasets for each proportions (see the detailed description of the scaling process in Methods Section 2.2). When focusing on the preservation of biological signals under label noise, scANVI ranks highest, with ssSTACAS showing comparable performance. Both methods perform similarly to the five unsupervised approaches, suggesting robustness to incorrect labels—likely due to their derivation from unsupervised frameworks, scVI and Seurat RPCA, respectively. As for the batch-effect removal performance, scDREAMER surpassed all other semi-supervised and unsupervised integration methods, outperforming the second best method, scCRAFT, by 6.83% with 70% of wrong labels. Notably, scDREAMER's top performance in batch correction task doesn't change much when wrong labels exist and when the proportion increases. scCRAFT and scANVI reach a balance between these two tasks, with both scores above the average (Figure S6b).

For semi-supervised methods, the performance of each method declines in bio-conservation task when the proportion of wrong labels increases. However, scANVI and ssSTACAS's performance in batch correction task increases as the proportion increases, while the other three methods show the opposite trend. Specifically, ItClust observes a significant drop in methods performance, especially in bio conservation task. From 30% to 50%, ItClust's performance observes a drop of 42.27% for bio conservation score and 19.22% for batch correction score after scaling, indicating its high sensitivity to cell labels' availability. ssSTACAS and scANVI are the most stable semi-supervised methods in this setting with nearly consistent performance across all proportions of missingness, similar to the performance when full labels are presented.

When comparing the performance of semi-supervised methods between settings with fully correct labels and those with mislabeled data, we observe that—with the exception of scDREAMER—most methods do not show significant improvement over unsupervised approaches in batch correction. Moreover, their performance in biological conservation degrades to varying degrees as the proportion of incorrect labels increases, in some cases falling well below that of unsupervised methods, despite previously outperforming them under clean label conditions.

From a closer view, we chose four datasets, human pancreas, macaque, human immune and lung atlas datasets as example, covering the relative easiest to hardest datasets, to comprehensively represent the performance of these methods under different scenario. Focusing on methods' performance on each metric (Figure S6c), we find that scDREAMER outperforms other methods in true positive rate and kBET Accept rate except for the macaque datasets, higher than the scCRAFT, which contributes to its superior performance in the batch integration task. Semi-supervised methods did not show significant improvement over unsupervised methods when it comes to other metrics.

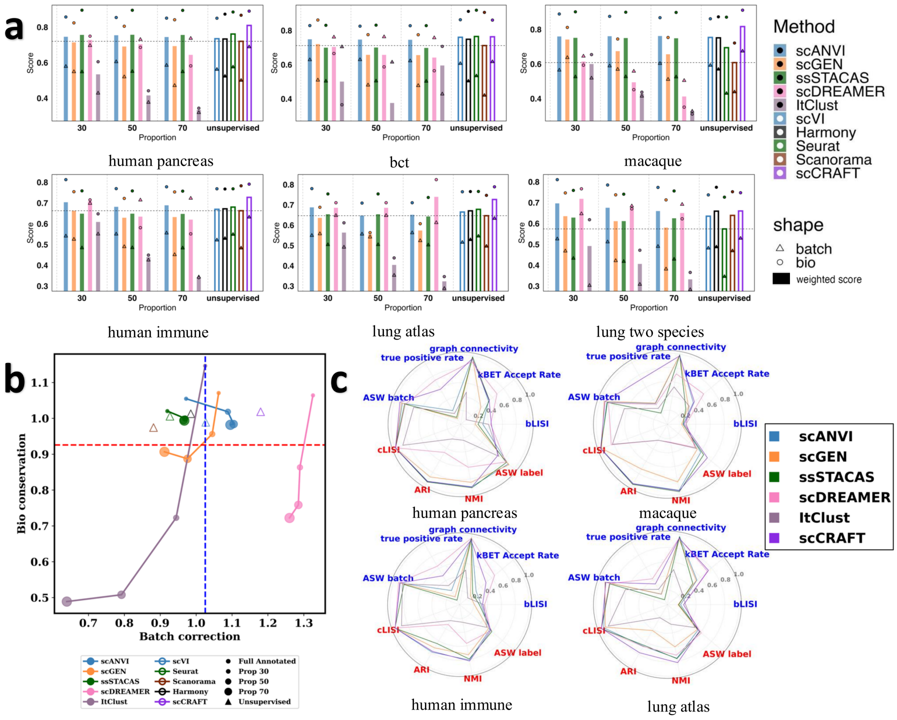

**Fig. S6** Partial Label Scenario II: Randomly Wrong Labels.
(**a**) Bar plots showing the performance of all methods across six datasets under this setting. Each bar represents the overall weighted score of a method; triangles and circles indicate the batch correction and biological conservation scores, respectively. The vertical dashed lines divide methods into four groups, namely, 30%, 50%, 70% and unsupervised approaches. The five unsupervised methods are shown on the right, represented by unfilled bars. (**b**) Scatter plot of the scaled batch correction score against the bio-conservation score for each method under the setting for different proportions, averaged across six datasets. The scaled score for each dataset and missing proportion is calculated as the ratio of overall bio-conservation/batch-mixing metric for a given method with respect to the corresponding mean using five unsupervised methods. The detailed scaling procedure can be found in Methods Section 2.2. Scaled scores for unsupervised methods are also included using unfilled triangles. Different colors indicate the methods and the size of dot shapes represent the missing proportions. The horizontal red dashed line represents the average bio conservation score across all methods (both supervised and unsupervised methods), while the vertical blue dotted line represented the average batch correction score. (**c**) Radar plots showing the performance of all methods on individual metrics for the *human pancreas, macaque, human immune,* and *lung atlas* datasets, averaged over all the three proportions for semi-supervised methods. Metrics include biological conservation (red) and batch correction (blue). As scCRAFT achieved the highest overall performance among unsupervised methods, only its scores are shown for clarity; radar plots for the remaining methods are provided in the Supplementary Note 5.

### 5 Scenario III: Harmony Integration Results of Each Datasets Under Different Proportion Thresholds

In the main text, we assign the missing and mixing label according to the Harmony clustering results for each datasets, and here we present the visualization of clustering results (See fig. Figure S7).

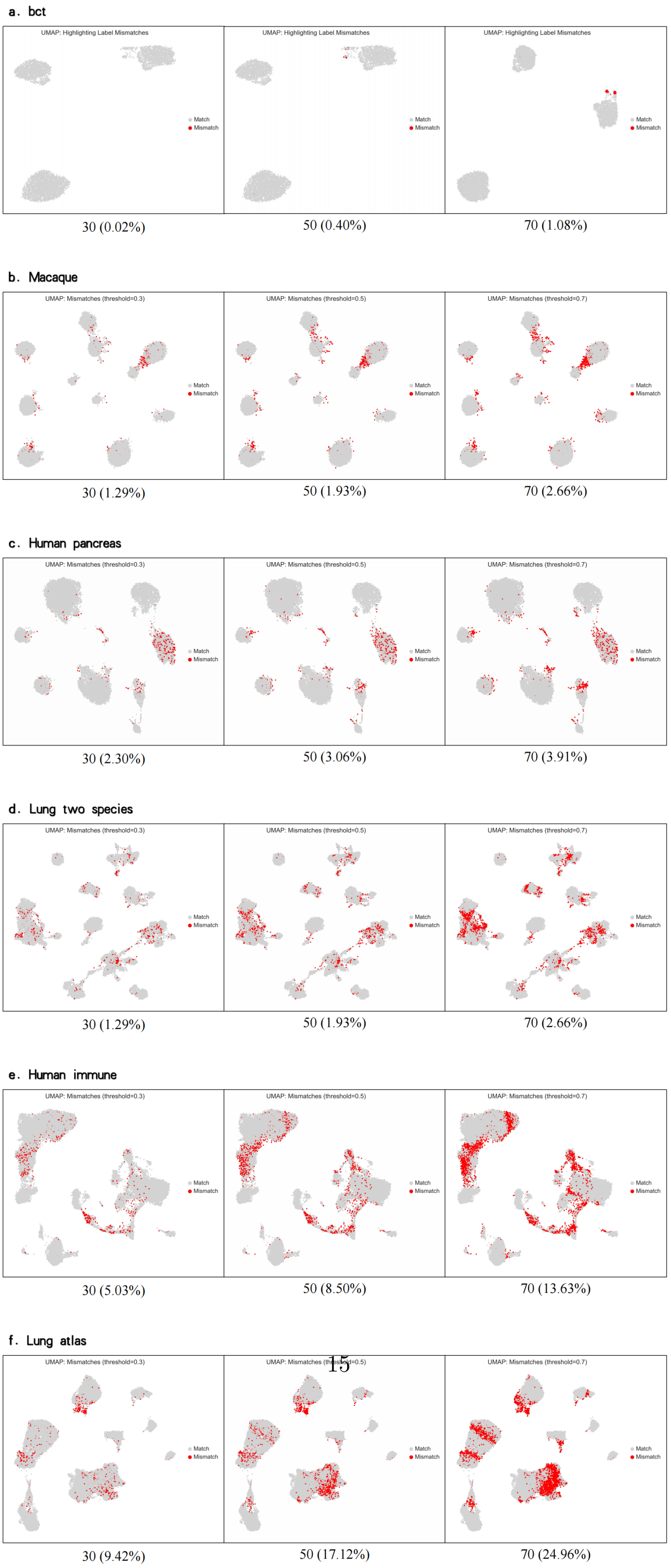

**Fig. S7** Harmony Clustering Results for each datasets in Missing and Mixing at Edge Setting
The plot shows Harmony clustering for five datasets (subpanels a–f: bct, macaque, human pancreas, lung two species, human immune, and lung atlas). In each subpanel, the three side-by-side plots correspond to the three predefined proportion thresholds. Within every plot, red cells mark missing or mixed assignments and gray cells mark correctly resolved cells. The label beneath each plot gives the proportion threshold value, with the mismatch fraction (mismatched cells/total cells) in parentheses.

The detailed reassigned proportion in each datasets at the threshold of $\gamma = 30\%$, $50\%$ and $70\%$ is reported in the table below. The proportion of cells whose labels were set to missing ("Unknown") is also reported in the parenthesis.

**Table S1** Proportion of Reassigned Cells

| **Dataset** | $\gamma = 30\%$ | $\gamma = 50\%$ | $\gamma = 70\%$ |
|---|---|---|---|
| BCT | 0.02 (0.00) | 0.40 (0.00) | 1.08 (0.00) |
| Macaque | 1.29 (0.22) | 1.93 (0.38) | 2.66 (0.88) |
| Human Pancreas | 2.30 (0.28) | 3.06 (0.55) | 3.91 (1.02) |
| Lung Two Species | 4.67 (0.55) | 8.12 (1.97) | 12.84 (4.61) |
| Human Immune | 5.03 (0.78) | 8.50 (2.10) | 13.63 (5.15) |
| Lung Atlas | 9.42 (1.32) | 17.12 (4.31) | 24.96 (8.84) |

### 6 Scenario IV: Partially Annotated Batches Results for Lung Two Species and BCT Datasets

Given that lung two-species and BCT datasets contain only two and three batches respectively (as noted in the main text), proportion-based selection of unannotated batches was impractical. Consequently, we manually designated one or two batches in each dataset as unannotated. ssSTACAS requires multiple annotated batches to operate in semi-supervised mode; therefore, we exclude it from evaluations on these two datasets. The corresponding results are presented in bar plots below. Semi-supervised methods generally underperform unsupervised methods, mirroring the trend observed in the other four datasets as discussed in the main text.

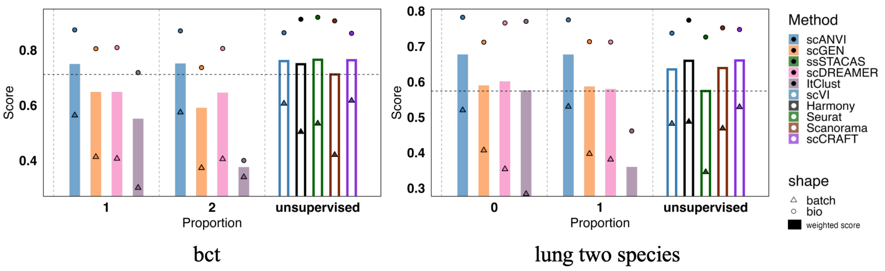

**Fig. S8** Bar plots for lung two species and bct datasets in Partially Annotated Batches Setting Bar plots showing the performance of all methods (ssSTACAS was excluded due to its design) across lung two species and bct datasets under this setting. Each bar represents the overall weighted score of a method; triangles and circles indicate the batch correction and biological conservation scores, respectively. The vertical dashed lines divide methods into three groups. Results for BCT (three batches) are presented with 1 or 2 unannotated batches. Results for lung two-species (two batches) are presented with either zero or one unannotated batch. The five unsupervised methods are shown on the right, represented by unfilled bars.

### 7 Scenario V: Semi-Supervised Integration Methods' Performance on Auto-annotation Labels After Fuzzy Matching

In the auto-annotated setting, predicted labels can differ from the original annotations, sometimes identifying different cell types or providing a more granular classification. This discrepancy can lead to an underestimation of the performance of semi-supervised methods. To mitigate this, we propose using "fuzzy matching." This approach involves mapping predicted labels to existing ones based on prior biological knowledge, allowing us to test whether the method's performance is robust to variations in cell type labels. A subsequent improvement in performance metrics would serve as strong evidence for this hypothesis. Such a result would underscore the utility of a fuzzy matching strategy and suggest that it is a crucial step for the robust and accurate evaluation of semi-supervised annotation methods.

We performed fuzzy matching on human immune datasets with Azimuth predicted labels. Azimuth predicted more kinds of cell types than the given labels, so we mapped the Azimuth labels to the given labels based on our prior knowledge of cell types (See table S2).

**Table S2** Mapping of Given Cell Types to Azimuth Predicted Cell Types

| **Given Cell Type** | **Azimuth Predicted Cell Type(s)** |
|---|---|
| CD4+ T cells | CD4 CTL, CD4 Naive, CD4 Proliferating, CD4 TCM, CD4 TEM |
| CD8+ T cells | CD8 Naive, CD8 TCM, CD8 TEM |
| CD10+ B cells | None |
| CD14+ Monocytes | CD14 Mono |
| CD16+ Monocytes | CD16 Mono |
| CD20+ B cells | B intermediate, B memory, B naive |
| Erythrocytes | Eryth |
| Erythroid progenitors | None |
| HSPCs | HSPC |
| Megakaryocyte progenitors | Platelet |
| Monocyte progenitors | None |
| Monocyte-derived dendritic cells | cDC1, cDC2 |
| NK cells | NK, NK Proliferating, NK_CD56bright |
| NKT cells | dnT |
| Plasma cells | Plasmablast |
| Plasmacytoid dendritic cells | pDC |

To assess the impact of label refinement, we reran the integration analysis using fuzzy matching labels in place of the original raw Azimuth labels. Overall, this change had a limited effect on most metric scores and the relative rankings of the methods for overall weighted scores, batch correction, and bio conservation (see Figure S9, Figure S10, and Table S3).

While most methods showed little change, ItClust experienced the most significant impact, with its ARI, graph connectivity, and true positive rate each increasing by roughly 0.1. Both scDREAMER and scGEN also saw moderate gains in true positive rate. However, scGEN's improvement was counterbalanced by declines in its kBET acceptance rate and bLISI. Notably, scGEN was the only method to register a drop in its overall weighted score after adopting fuzzy matching labels; the other four methods all improved (Figure S9).

The ranking between methods saw minor shifts: scANVI moved ahead of scGEN in the overall weighted score, and ItClust surpassed scGEN in the bio conservation score. Other rankings were unaffected (Table S3).

In conclusion, the performance of the integration methods remained largely stable despite the change to fuzzy matching labels. This suggests that utilizing the initial raw predicted auto-annotation labels is a viable and sound approach for this integration.

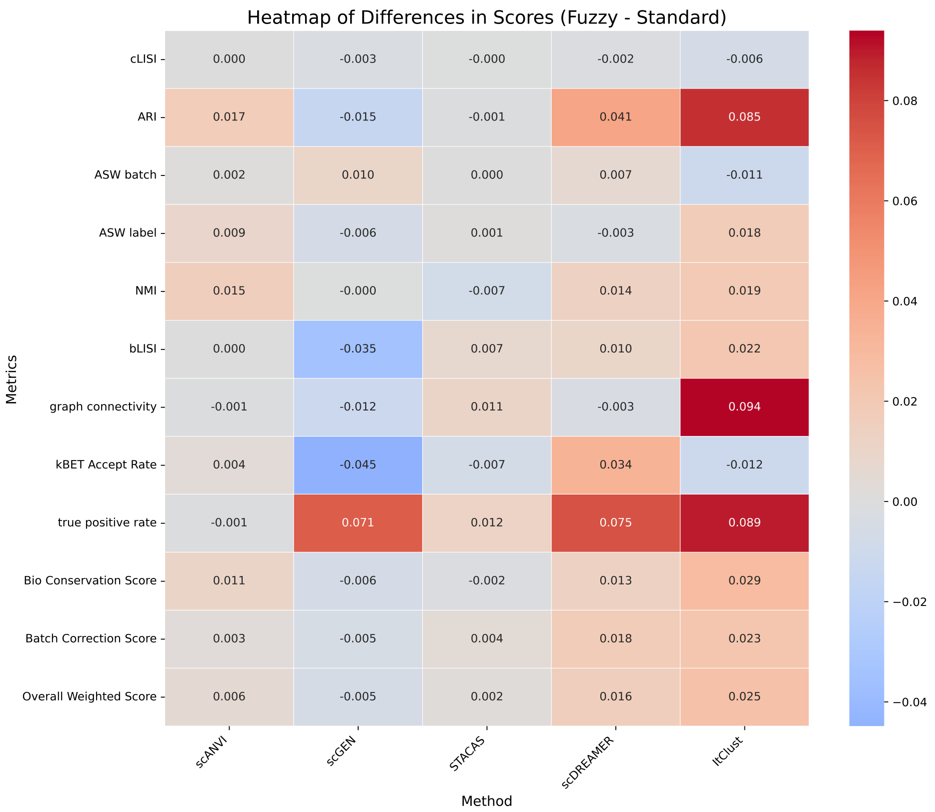

**Fig. S9** Heatmap for Difference between Fuzzy Matching Labels and Raw Azimuth Labels
This plot presents the differences in a grid format. Rows represent the Methods, and columns represent metrics. The color intensity and hue of each cell directly visualize the calculated difference. The plot provides a quick overview of patterns in the differences across all methods and scores simultaneously, where we can easily spot clusters of large positive or negative differences.

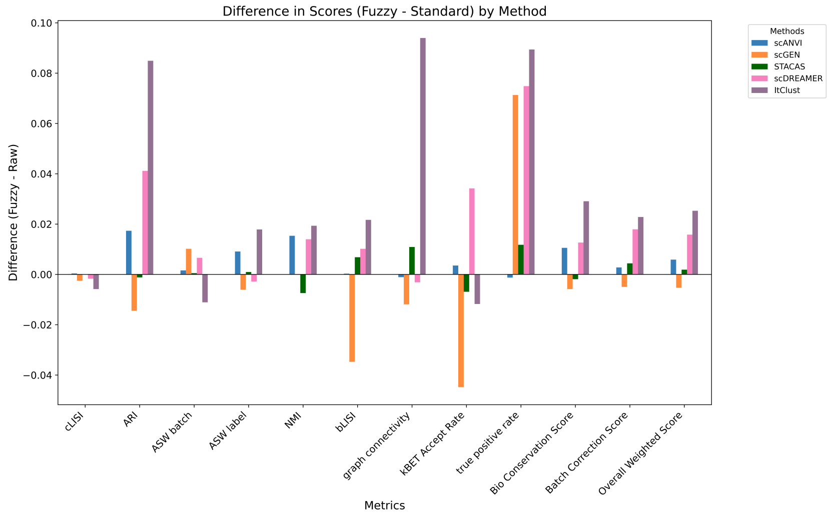

**Fig. S10** Bar Chart for Difference between Fuzzy Matching Labels and Raw Azimuth Labels
For each 'Metrics', this chart displays a group of bars. Each bar in the group corresponds to one of the methods. The height of the bar indicates the difference in scores between the two files for that specific method and score type. The plot precisely compares the magnitude and direction (positive or negative) of differences for each metric across all methods, where we can easily see which methods have larger or smaller changes in specific scores.

**Table S3** Rank Comparisons for Bio Conservation, Batch Correction, and Overall Weighted Scores between Using Raw Azimuth Predicted Labels and Fuzzy Matching Azimuth Labels

| **Method** | **Bio Conservation** | | **Batch Correction** | | **Overall Weighted** | |
|---|---|---|---|---|---|---|
| | **Fuzzy** | **Raw** | **Fuzzy** | **Raw** | **Fuzzy** | **Raw** |
| scANVI | 1 | 1 | 4 | 4 | 3 | 4 |
| scGEN | 5 | 4 | 2 | 2 | 4 | 3 |
| ssSTACAS | 2 | 2 | 3 | 3 | 2 | 2 |
| scDREAMER | 3 | 3 | 1 | 1 | 1 | 1 |
| ItClust | 4 | 5 | 5 | 5 | 5 | 5 |
